# Control iPSC lines with clinically annotated genetic variants for versatile multi-lineage differentiation

**DOI:** 10.1101/666560

**Authors:** Matthew R Hildebrandt, Miriam S Reuter, Wei Wei, Naeimeh Tayebi, Jiajie Liu, Sazia Sharmin, Jaap Mulder, L Stephen Lesperance, Patrick M Brauer, Caroline Kinnear, Alina Piekna, Asli Romm, Jennifer Howe, Peter Pasceri, Rebecca S Mok, Guoliang Meng, Matthew Rozycki, Deivid de Carvalho Rodrigues, Elisa C Martinez, Michael J Szego, Juan Carlos Zúñiga-Pflücker, Michele K Anderson, Steven A Prescott, Norman D Rosenblum, Binita M Kamath, Seema Mital, Stephen W Scherer, James Ellis

**Affiliations:** Developmental and Stem Cell Biology, The Hospital for Sick Children, Toronto ON, Canada; Genetics and Genome Biology, The Hospital for Sick Children, Toronto, ON, Canada; The Centre for Applied Genomics, The Hospital for Sick Children, Toronto, ON, Canada; Neurosciences and Mental Health, The Hospital for Sick Children, Toronto, ON, Canada; Department of Immunology, University of Toronto, Sunnybrook Research Institute, Toronto, ON, Canada; Department of Molecular Genetics, University of Toronto, Toronto, ON, Canada; Dalla Lana School of Public Health, University of Toronto, Toronto, ON, Canada; Department of Family and Community Medicine, University of Toronto, Toronto, ON, Canada; The Joint Centre for Bioethics, University of Toronto, Toronto, ON, Canada; Unity Health Toronto, Toronto, ON, Canada; Institute of Biomaterials and Biomedical Engineering, University of Toronto, Toronto, ON, Canada; Department of Physiology, University of Toronto, Toronto, ON, Canada; Department of Pediatrics, University of Toronto, Toronto, ON, Canada; McLaughlin Centre, University of Toronto, Toronto, ON, Canada

## Abstract

Induced Pluripotent Stem Cells (iPSC) derived from healthy individuals are important controls for disease modeling studies. To create a resource of genetically annotated iPSCs, we reprogrammed footprint-free lines from four volunteers in the Personal Genome Project Canada (PGPC). Multilineage directed differentiation efficiently produced functional cortical neurons, cardiomyocytes and hepatocytes. Pilot users further demonstrated line versatility by generating kidney organoids, T-lymphocytes and sensory neurons. A frameshift knockout was introduced into *MYBPC3* and these cardiomyocytes exhibited the expected hypertrophic phenotype. Whole genome sequencing (WGS) based annotation of PGPC lines revealed on average 20 coding variants. Importantly, nearly all annotated PGPC and HipSci lines harboured at least one pre-existing or acquired variant with cardiac, neurological or other disease associations. Overall, PGPC lines were efficiently differentiated by multiple users into cell types found in six tissues for disease modeling, and clinical annotation highlighted variant-preferred lines for use as unaffected controls in specific disease settings.

## INTRODUCTION

The development of induced pluripotent stem cells (iPSC) (Takahashi et al., 2007; Yu et al., 2007) led to rapid development of many stem cell-based models of disease (Dimos et al., 2008; Park et al., 2008). Despite exponential growth in the application of iPSCs across multiple tissue and organ-based systems, there remains no consistent consensus about which control lines should be used in disease modeling studies. Over the past decade, choices for control cells have ranged from: 1) human embryonic stem cells (hESCs) that are considered healthy despite a medical history being unavailable (Dimos et al., 2008; Marei et al., 2017; Park et al., 2008), 2) iPSCs from healthy but unrelated individuals (Fernandes et al., 2016; Schwartzentruber et al., 2018), 3) iPSCs from unaffected family members who may have been phenotyped for the disease of interest, but with unknown broader health profile (Lan et al., 2013), and 4) isogenic pairs of iPSC lines derived through CRISPR-Cas9 gene editing (Deneault et al., 2018; Li et al., 2013; Ma et al., 2018; Mosqueira et al., 2018; Wang et al., 2018), or through non-random X chromosome inactivation status in female cells (Cheung et al., 2011; Pomp et al., 2011; Tchieu et al., 2010). Hundreds of sources of unrelated and related healthy iPSC lines exist and are widely available from individual labs, biobanks and large iPSC-focused consortia such as: HipSci, IPSCORE, Progenitor Cell Biology Consortium and NextGen (D’Antonio et al., 2017; Panopoulos et al., 2017; Salomonis et al., 2016; Streeter et al., 2017).

Although there are genetically diverse lines to reflect heterogeneity found within the human population, all control lines are potentially compromised by genetic variants that may predispose to a phenotype or mask it (DeBoever et al., 2017; Hollingsworth et al., 2017). To date, disease modeling has focused on penetrant monogenic disorders that may be relatively unaffected by the presence of concurrent variants (Cheung et al., 2011; Lan et al., 2013; Li et al., 2013). However, we anticipate an emerging need for healthy controls with few disease variants as modeling of complex diseases builds towards assessing the impact of modifier genes or multigenic disorders that may involve multiple variants as well as noncoding variants in gene regulatory regions.

iPSCs carry additional variants compared to donor sequences (D’Antonio et al., 2018; Gore et al., 2011). This has made apparent the need for whole genome sequencing (WGS) in order to identify the full set of potential disease-susceptibility variants present in such control lines (Bhutani et al., 2016; Burrows et al., 2016; D’Antonio et al., 2018; Gore et al., 2011; Kilpinen et al., 2017; Popp et al., 2018). Although there are some common reprogramming-associated variants (Yoshihara et al., 2017), most variants appear to be present in the original mosaic source of cells reprogrammed (Abyzov et al., 2017; Young et al., 2012). Some of these variants could affect downstream differentiations and baseline phenotypes of differentiated lineages (Hoekstra et al., 2017). Further, most control lines are recruited for specific studies limited to a single tissue type or disease, and therefore their versatility for multi-lineage directed differentiation into many functional cell types required for broad disease modeling research is not firmly established.

One way to limit the presence of potentially confounding variants is to reprogram cells from selected donors who have minimal variant load. In both the initial Personal Genome Project (PGP) and Personal Genome Project Canada (PGPC) publications, one aim was to generate iPSCs that would have extensive genomic characterization (Ball et al., 2012; Reuter et al., 2018). PGPC genotyped and clinically annotated the genomes of 56 apparently healthy individuals who consented to disclosure of their genome sequence and medical traits (Reuter et al., 2018). In addition to comprehensive annotation of all classes of constitutional genetic variants, these analyses also included their assessment of the mitochondrial genomes and their pharmacogenetic diplotypes. All healthy PGPC individuals harbour heterozygous variants of unknown significance in disease relevant genes, but still had no overt disease phenotype at the time of initial assessment or at the start of this study. Here we report the iPSC resource generated from PGPC donors.

Our resource comprises multiple iPSC lines derived from two male and two female donors. One line each from both males and one female was subjected to multi-lineage directed differentiation into cortical neurons, cardiomyocytes and hepatocytes representative of the three germ layers. The morphology and function of the resulting cells were evaluated to assess the versatility of PGPC iPSC lines for *in vitro* studies of different tissues. To further evaluate the versatility of the resource, we shared the three-best characterized PGPC lines with pilot users for differentiation into kidney organoids, T-lymphocytes, and sensory neurons. CRISPR gene editing of a known cardiomyopathy gene created an isogenic pair of lines for modeling a cardiac disorder. As variant annotation of the donors became available (Reuter et al., 2018), we performed WGS to search for iPSC line-specific variants that were distinct from donor PGPC blood variants, and surveyed off-target mutations in the gene edited line.

## RESULTS

### Isolation and pluripotency characterization of PGPC iPSC lines

We invited PGPC donors to participate in this iPSC study, and selected two male (PGPC3 and PGPC17) and two female donors (PGPC14 and PGPC1) (Reuter et al., 2018). We collected peripheral blood to isolate and reprogram CD34+ cells using non-integrating Sendai viruses. Approximately 120 clones from each donor were picked and qualitative metrics (colony morphology and low levels of spontaneously differentiated cells) were used to select lines for characterization. iPSC lines were maintained in feeder-free conditions and tested for Sendai virus clearance at passage (P)8 to 10. Sendai virus negative lines were sent for karyotyping between P13-15. At least four karyotypically normal cell lines were found from each donor with standard characterization results summarized in Supplemental Table 1 and representative data shown in Fig. S1. All cell lines stained positive for both cytoplasmic (SSEA4 and TRA-1-60) and nuclear (POUF51 and NANOG) pluripotency markers (Fig. S1). We tested functional pluripotency by spontaneously differentiating embryoid bodies followed by staining for markers of all three germ layers—ectoderm (TUBB3), mesoderm (SMA), and endoderm (AFP) (Fig. S1). All female lines had skewed X chromosome inactivation as revealed by androgen receptor assays consistent with preservation of an inactive X chromosome observed in isogenic female lines (Fig. S1). These data confirm basic pluripotency status of our resource and cells were expanded and banked at passages ranging from P14-16.

**Table 1.** Reprogrammed PGPC iPSC lines: Novel genomic variants (overview).

| PGPC3_75 | SNVs/indels |  |  |  | CNVs |  |
| --- | --- | --- | --- | --- | --- | --- |
|  | All | Exonic | Non-synonymous | Loss-of-function | All | Exonic |
| Genome-wide | 1981 |  |  |  | 0 |  |
| All genes | 847 | 19 | 12 | 1 | 0 | 0 |
| OMIM genes | 172 | 3 | 3 | 0 | 0 | 0 |
| Constrained genes* | 216 | 6 | 5 | 0 | 0 | 0 |
| PGPC14_26 | SNVs/indels |  |  |  | CNVs |  |
|  | All | Exonic | Non-synonymous | Loss-of-function | All | Exonic |
| Genome-wide | 1235 |  |  |  | 1 |  |
| All genes | 499 | 18 | 13 | 2 | 1 | 0 |
| OMIM genes | 104 | 3 | 2 | 0 | 1 | 0 |
| Constrained genes* | 150 | 8 | 8 | 2 | 1 | 0 |
| PGPC17_11 | SNVs/indels |  |  |  | CNVs |  |
|  | All | Exonic | Non-synonymous | Loss-of-function | All | Exonic |
| Genome-wide | 1169 |  |  |  | 0 |  |
| All genes | 466 | 23 | 14 | 1 | 0 | 0 |
| OMIM genes | 95 | 6 | 6 | 1 | 0 | 0 |
| Constrained genes* | 113 | 5 | 5 | 1 | 0 | 0 |
| PGPC17_11<br>MYBPC3_KO | SNVs/indels |  |  |  | CNVs |  |
|  | All | Exonic | Non-synonymous | Loss-of-function | All | Exonic |
| Genome-wide | 917 |  |  |  | 1 |  |
| All genes | 382 | 17 | 9 | 1 | 0 | 0 |
| OMIM genes | 85 | 4 | 2 | 0 | 0 | 0 |
| Constrained genes* | 35 | 7 | 4 | 0 | 0 | 0 |

\* pLI>0.9 (http://exac.broadinstitute.org/).
3 PGPC3\_75, PGPC14\_26, and PGPC17\_11 were compared to the sequence data 4 obtained from whole blood. PGPC17\_11 MYBPC3\_KO was compared to the 5 PGPC17\_11 reprogrammed line. Abbreviations: CNV, copy number variant; indel, 6 insertion/deletion; SNV, single nucleotide variant.

We chose to focus on one cell line from the first three donors for deeper characterization as PGPC1 was recruited much later. PGPC3_75, PGPC14_26, and PGPC17_11 were selected for further phenotyping based on qualitative metrics regarding their growth rate, morphology, and relative low rate of spontaneous differentiation. RNA sequencing was analyzed online using Pluritest [pluritest.org; (Müller et al., 2011)] and all three lines cluster to the pluripotency quadrant (Fig. S1). As explained in detail below, we validated the pluripotency and explored the versatility of all three lines for multi-lineage directed differentiation to excitatory cortical neurons, cardiomyocytes, and hepatocytes as representatives of cells derived from ectoderm, mesoderm and endoderm respectively.

At this point the WGS data of all the PGPC participants became available and were annotated for coding variants defined by the American College of Medical Genetics (ACMG) (Richards et al., 2015). Two heterozygous variants of uncertain clinical significance (VUS) associated with electrophysiological alterations in cardiac disease (Table S2) were identified in PGPC3 [TRPM4 (Liu et al., 2010)] and PGPC14 [KCNE2 (Gordon et al., 2008)], respectively. Additional VUSs that could impact neurologic function were found in cells derived from PGPC14 and PGPC17 (Table S3). We therefore prioritized PGPC3 as a preferred line for neuronal models and PGPC17 as a variant-preferred line for cardiac models based on their pre-existing variants. The newest PGPC1 female lines are cardiac variant-preferred and are available only with donor variant annotation (Table S2) and pluripotency characterization as part of the resource.

### Ectodermal differentiation into active cortical neurons

To evaluate PGPC iPSC-derived neurons, we infected PGPC lines and a previously published control iPSC line (WT37) (Cheung et al., 2011) with lentivirus bearing doxycycline-inducible *NEUROG2* to generate homogenous populations of excitatory cortical neurons (Zhang et al., 2013). Neurons were induced with doxycycline for one week and selected with puromycin and Ara-C (Deneault et al., 2018, 2019) in 6-well plates then re-seeded to 24-well dishes for morphological analysis in co-cultures with mouse astrocytes over seven weeks (Fig. 1A). To measure single neurons, we sparsely labeled six-week cultured neurons by transfection with ubiquitous expressing GFP plasmid in 2 batches. Neurons were identified by staining with pan-neuronal marker MAP2 (Fig. 1B). Soma area, dendritic length and neuronal complexity of the PGPC neurons determined by Sholl analysis were similar to the WT control (Fig. 1C-E).

**Figure 1.**
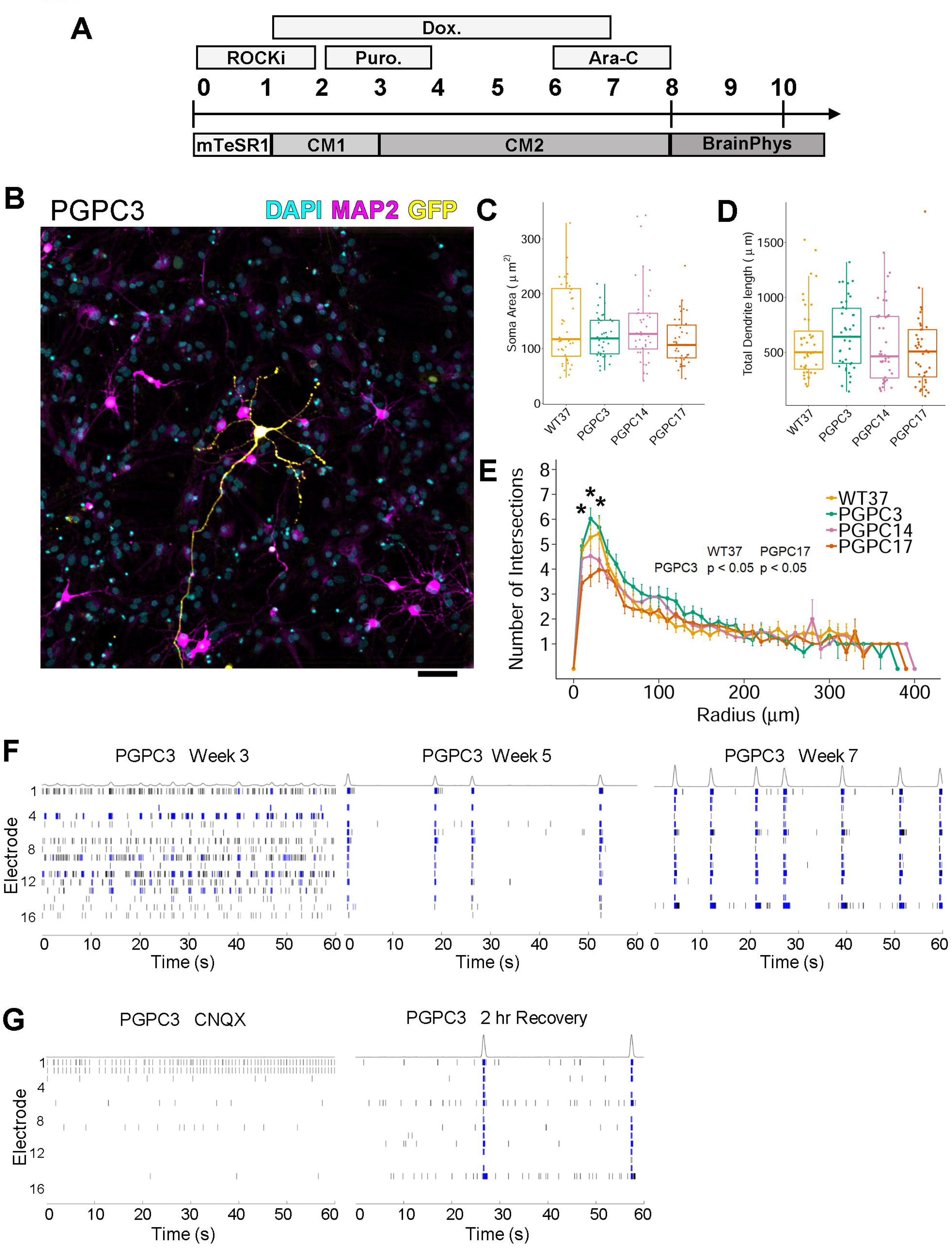
Active neurons generated from PGPC iPSCs display similar dendrite morphology and network circuitry. (A) Differentiation scheme to generate excitatory cortical neurons by induction of NGN2. Transduced iPSCs were dissociated to single cells and plated in the presence of ROCK inhibitor. At D1, media was changed to CM1 and NGN2 was induced by incubating with doxycycline until D7. Puromycin was added from D2-4 to remove any non-transduced cells. Culture media was changed to CM2 on D3. Ara-C was added from D6-8 to remove any remaining dividing cells. On D8, neurons are re-seeded for downstream assays in BrainPhys media. (B) Representative immunocytochemistry image of iPSC-derived neurons after six weeks in culture labeled with DAPI and MAP2 and sparsely labeled with GFP (biological replicates = 2; technical replicates = 20). Scale bar represents 100 *µ*m). (C-E) Plots of (C) soma area (D) total dendrite length (E) number of intersections (Sholl analysis) from six-week old neurons (biological replicates n = 2; technical replicates per batch = 20). (C-D) Box plots indicate median values. (E) Mean values were plotted with error bars indicating standard error. Statistically significant pair-wise comparisons indicated by * are inset. (F-G) Representative raster plots of PGPC3 neurons from a single well of recordings collected by micro-electrode arrays at different time points (two biological replicates each with 8 technical replicates). Each spike is indicated by a black line, blue lines represent bursts defined by at least five spikes each separated by an inter-spike interval of no more than 100 ms. (G) Bursts are absent after treatment of week 7 neurons with CNQX (left). Burst begin to return after compound removal and replacement with fresh basal media (right).

To investigate activity of the variant-preferred PGPC3 neurons, we collected weekly recordings from 48-well MEA plates (Axion BioSystems) for extracellular electrophysiology measurements (Deneault et al., 2018, 2019) over 6 weeks (weeks 2-7). Representative raster plots of PGPC3_75 showed progression of spontaneous activity at three weeks compared to development of network bursts at 5 and 7 weeks (Fig. 1F). At the 7 seven-week time point, we observed synchronous firing across multiple electrodes (minimum 8/16 electrodes) within wells, indicative of neural circuit formation as measured by network burst frequencies. Neurons displayed weighted Mean Firing Rates (wMFR) ranging from 5 to 7.5 Hz and network burst frequencies ranging from 0.1 to 0.35 Hz, which were comparable to or more active (∼5 Hz and ∼0.1 Hz respectively) than our previously published MEA results from NGN2-derived neurons (Deneault et al., 2018, 2019). To confirm that recorded activity was due to synaptic transmission from glutamatergic excitatory neurons, we treated cells with an α-amino-3-hydroxy-5-methyl-4-isoxazolepropionic acid (AMPA) receptor inhibitor—6-cyano-7-nitroquinoxaline-2,3-dione (CNQX)—which abolished network bursting (Fig. 1G). Network bursting began to recover two hours after washing out CNQX. These findings demonstrate differentiation of three PGPC lines into sparsely labeled neurons and that the variant-preferred PGPC3 line was spontaneously active in network circuits.

### Mesodermal differentiation into contractile cardiomyocytes

PGPC iPSCs were differentiated into cardiomyocytes (CMs) using a STEMdiff Cardiomyocyte Differentiation Kit (Fig. 2A). We observed beating cells at day 8 with all lines. Contractile cultures were dissociated to single CMs at D16 for flow cytometry. The proportion of cardiac troponin T (cTNT) positive cells was routinely between 75-85% (Fig. 2B). The D16 CMs were re-seeded into 24-well plates and matured for an additional 17 days to D33. Immunostaining showed that D33 CMs were a mixture of round and cylindrical shaped cardiomyocytes and most cells positively stained for cTNT, myosin light chain variant 2 (MLC2V—a ventricular marker), and the sarcomere marker alpha-actinin (Fig. 2C).

**Figure 2.**
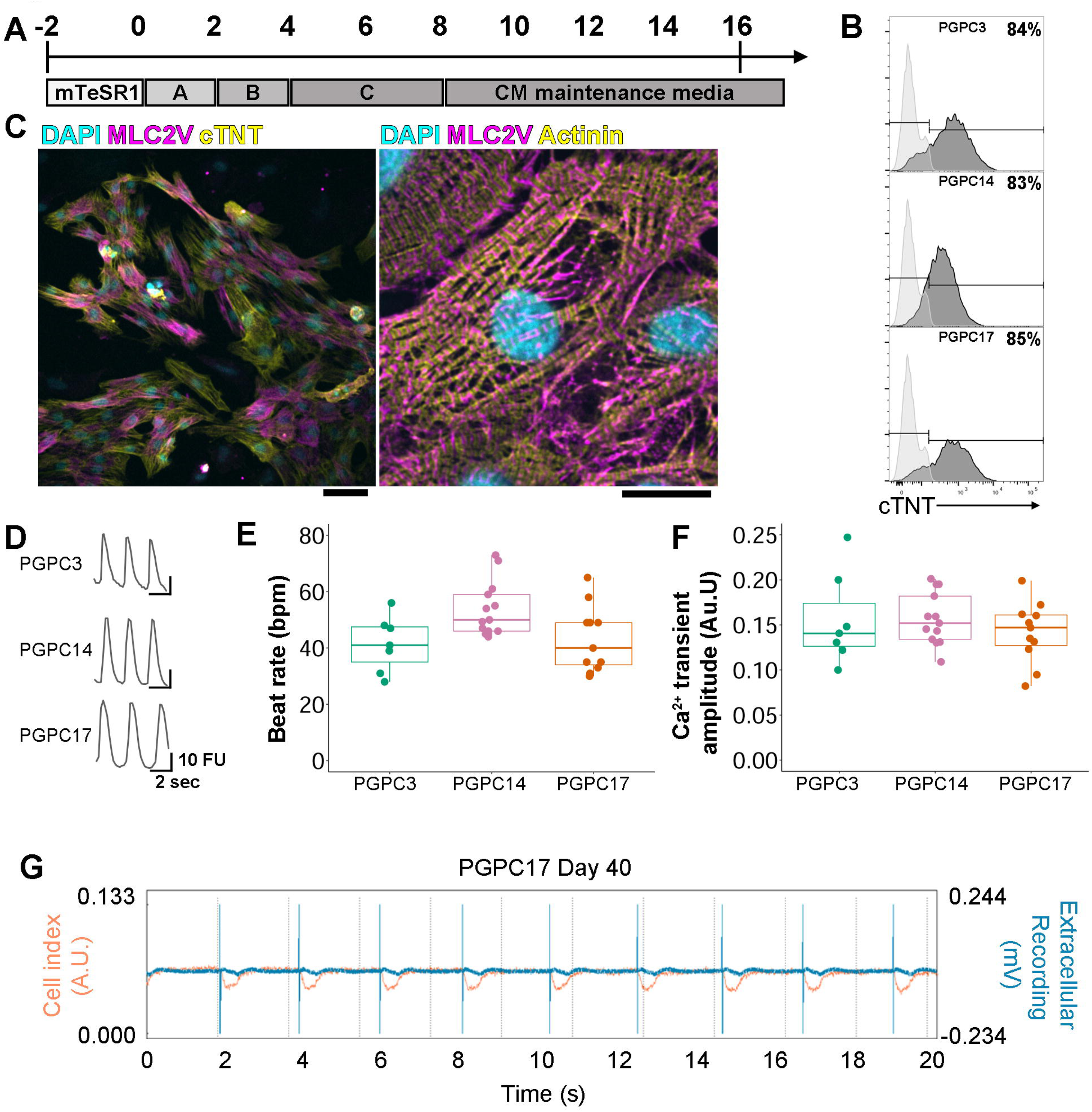
PGPC iPSCs differentiate to beating contractile cardiomyocytes. (A) Differentiation scheme to generate CMs using STEMdiff cardiomyocyte differentiation kit. iPSCs were dissociated to single cells, plated in 12-well plates, and allowed to reach 85-90% confluency before beginning differentiation. (B) D16 CMs were dissociated to single cells for reseeding and a proportion was labeled with anti-cTNT-FITC and subjected to flow cytometry (biological replicates ≥ 3). (C) Representative images of immunocytochemistry staining of D30 PGPC17 CMs labeled with DAPI, anti-MLC2V (both), and anti-cTNT (left) or anti-alpha actinin (right) (biological replicates = 2). Scale bars represent 100 *µ*m. (D) Representative traces of spontaneous Ca^2+^ transients of PGPC CMs at D31 measured by relative fluorescence intensity (biological replicates n ≥ 3; technical replicates per batch ≥ 2). (E-F) Plots of (E) beat rate and (F) Ca^2+^ transient amplitudes. (G) Representative xCELLigence data of D40 PGPC17 CMs showing impedance changes (BAmp: defined as the cell index value between lowest and highest points within a beat waveform) reflecting CM beat waveform and absolute extracellular voltage tracings over a 20 second recording (biological replicates = 3; technical replicates ≥ 3).

For Intracellular Ca^2+^ transient measurements in 96 well plates between D31 and D34, the CMs were loaded with 1 *µ*M Fluo-4 AM dye (Invitrogen) for 30 min at 37°C. Fluorescence intensity ratios were plotted against time to calculate the Ca^2+^ transient amplitude and rate (Fig. 2D-F). All three PGPC-CMs had similar average beat rates and amplitudes. To measure contractility of PGPC17_11 in a complementary method and to determine extracellular electrophysiology, an xCELLigence Real Time Cell Analysis (RTCA) Cardio ExtraCellular Recording (ECR) system was used. In brief, contracting CMs in 48-well plates were recorded over 20 second sweeps every three hours for ∼25 days after reseeding (Fig. 2G). Contractility of CMs was evaluated via impedance readouts as beats per minute (bpm) and beating amplitude (BAmp) defined as the cell index value between lowest and highest points within a beat waveform. Beat rate averaged 36 bpm (range 32 to 49 bpm) with average amplitude 0.04 arbitrary units (AU) (range 0.027 to 0.05 AU). Extracellular field potential spike amplitudes defined as the difference between the lowest and highest recorded voltages ranged from 0.12 to 0.55 mV. These experiments demonstrate differentiation of three PGPC lines into beating cardiomyocytes and highlight the potential value of using PGPC17 for CRISPR gene editing for cardiac disease modeling.

### Endodermal differentiation into enzymatically active hepatocytes

For endodermal differentiation we generated hepatocyte-like cells (HLC) using a protocol adapted from (Ogawa et al., 2015) (Fig. 3A). Differentiated cells were characterized at multiple stages to monitor quality and efficiency. At D4, over 95% of cells co-expressed definitive endoderm (DE) markers *CXCR4* and *cKIT* (data not shown). DE cells were induced to generate foregut (FG) progenitors as indicated with the increase in FG markers *FOXA2* and *GATA6* (normalized to iPSCs) compared to DE (Fig. 3B). FG progenitors were further specified to hepatoblasts (HBs) followed by maturation to HLCs by D25 where clear upregulation of respective mRNAs was assessed by qPCR (normalized to fetal liver) (Fig. 3B). Over 95% of HLCs tested positive via flow cytometry for hepatocyte markers including albumin (ALB), alpha fetoprotein (AFP), alpha-1-antitrypsin (A1AT), CYP3A7 (Fig. 3C) and further supported by immunostaining for AFP, ALB, CYP3A7, and HNF4A (Fig. 3D). Measuring functional activity of HBs (D14) and HLCs (D25) was performed using a p450-glo assay (Promega). As expected, HLCs had significantly more enzymatic activity of both CYP3A4 and CYP3A7 as measured by luminescence as compared to HBs. Treatment with 1 *µ*M ketoconazole inhibited enzymatic activity of CYP3A7 to levels observed in HBs (Fig. 3E). These results demonstrate differentiation of the 3 PGPC lines into hepatocytes that produce active enzymes.

**Figure 3.**
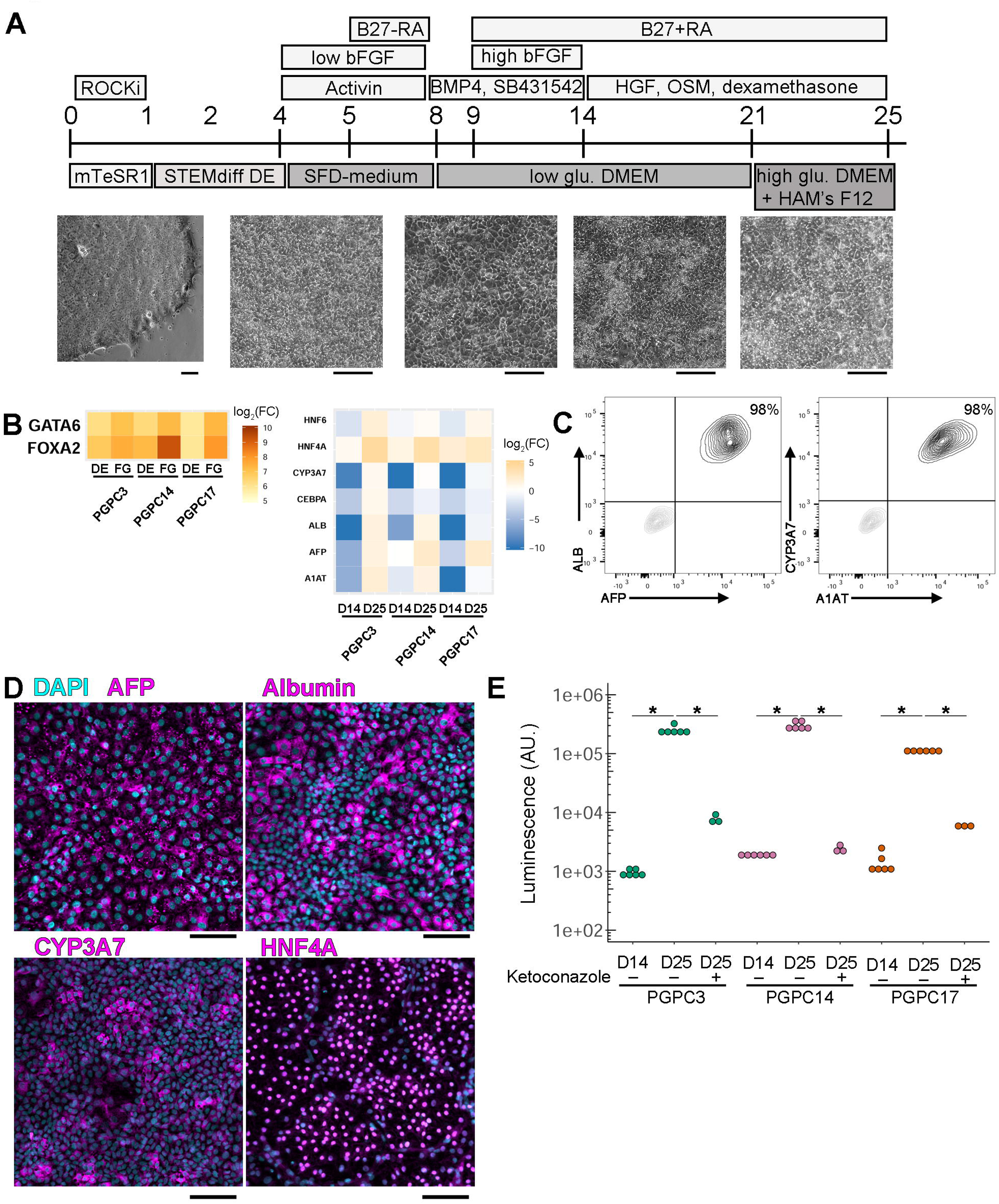
Enzymatically active HLCs are generated from PGPC iPSCs. (A) Hepatocyte-like cell differentiation scheme. iPSCs were dissociated to single cells and maintained in ROCK inhibitor for 24 hours to support survival. From D1-4 cells are transferred to STEMdiff Definitive Endoderm (DE) differentiation kit. From D4-8 cells were switched to serum-free-differentiation (SFD)-based medium with basic fibroblast growth factor (bFGF) and activin A for 4 days adding B27 without retinoic acid (RA) for days 5-8. Media was changed every other day, then on D8, cells were switched to low glucose DMEM supplemented with SB-431542 and bone morphogenic protein (BMP4) from day 8 to day 14. bFGF was added back to the media during D9-14 while B27 with retinoic acid was supplemented from D9-25 with media changed every other day. From D14-25 hepatocyte growth factor (HGF), dexamethasone and oncostatin M (OSM) were added to culture media. At D21 cells were cultured in a mixture of low glucose DMEM / Ham’s F12 (3:1) media. Bright field images highlight morphology changes during differentiation. Scale bars represent 100 *µ*m. (B) Heatmaps indicating log_2_ fold change of marker gene expression normalized to iPSCs (left) or to fetal liver (right) (biological replicates = 3). (C) Dissociated D25 PGPC14 HLCs were labeled with anti-ALB and□anti-AFP (left) or anti-alpha-1-AT and□ anti-CYP3A7 (right) and subjected to flow cytometry (biological replicates = 3). (D) Representative immunocytochemistry images of D25 PGPC14 HLCs labeled with DAPI and anti-AFP, anti-Albumin, anti-CYP3A7, or anti-HNF4A (biological replicates = 3). Scale bar represents 100 *µ*m. (E) Log scale plot of luminescence of each cell line measuring P450 enzymatic activity assayed at D14 and D25 with or without ketoconazole as an inhibitor (biological replicates n = 3; technical replicates for untreated samples = 2, treated samples = 1). Statistical significance was determined by Dunn’s test between all samples and * indicate pairs where p < 0.05.

### Utility of the resource – Mesodermal differentiation into kidney organoids and T-cells

To test the utility of PGPC lines as a resource, we made them available to pilot users. Unlike mono-layer differentiations described above, human kidney organoids are 3D structures generated from iPSCs consisting of multiple cell types and resembling early embryonic human kidney tissue (Morizane et al., 2015; Taguchi et al., 2014; Takasato et al., 2015; Wu et al., 2018). Kidney organoids were made from all three PGPC lines according to the protocol described by Takasato et al., 2015 but with minor modifications (e.g. feeder-free iPSC culture and omission of conditioned medium) similar to those implemented in more recent publications (van den Berg et al., 2018; Forbes et al., 2018) (Fig. S2A).

This protocol entailed a 7-day monolayer culture with directed differentiation towards posterior streak mesoderm (PSM) and subsequently to anterior and posterior intermediate mesoderm (respectively AIM and PIM). This was accomplished by applying the canonical WNT-signaling activator, CHIR99021 (CHIR), followed by a switch to fibroblast growth factor-9 (FGF9) and heparin. Timing of the FGF9/heparin switch (between D3-5 of differentiation) determined the relative proportion of AIM vs. PIM and thus, fewer or more nephrons (Takasato et al., 2015). For all experiments, we made this factor switch on D5. During this course, the PSM marker *T* (brachyury) was transiently induced followed by AIM marker *GATA3* and PIM marker *HOXD11* as measured by qPCR and expression was normalized to iPSCs at D0 (Fig. 4A). After seven days of monolayer differentiation, cells were aggregated by centrifugation and transferred to Transwell membranes. Aggregates were pulsed with CHIR for one hour and then treated with FGF9/heparin for five additional days. Shortly after aggregation, cellular reorganization began and resulted in formation of nephron-like structures in aggregates. mRNA level analysis (normalized to D7 pre-aggregated cells) showed the induction of markers of different nephron segments, endothelial, and stromal cell markers in D25 organoids (Fig. 4A). Immunofluorescence imaging of D18 cross-sections of organoids showed glomerular structures—positive for podocyte marker Wilms Tumour 1 (WT1)—as well as tubular structures, both proximal—labeled with lectin (LTL)—and distal—positive for E-Cadherin (ECAD) (Fig. 4B). These results show 3D kidney organoid structures are produced by the 3 PGPC lines.

**Figure 4.**
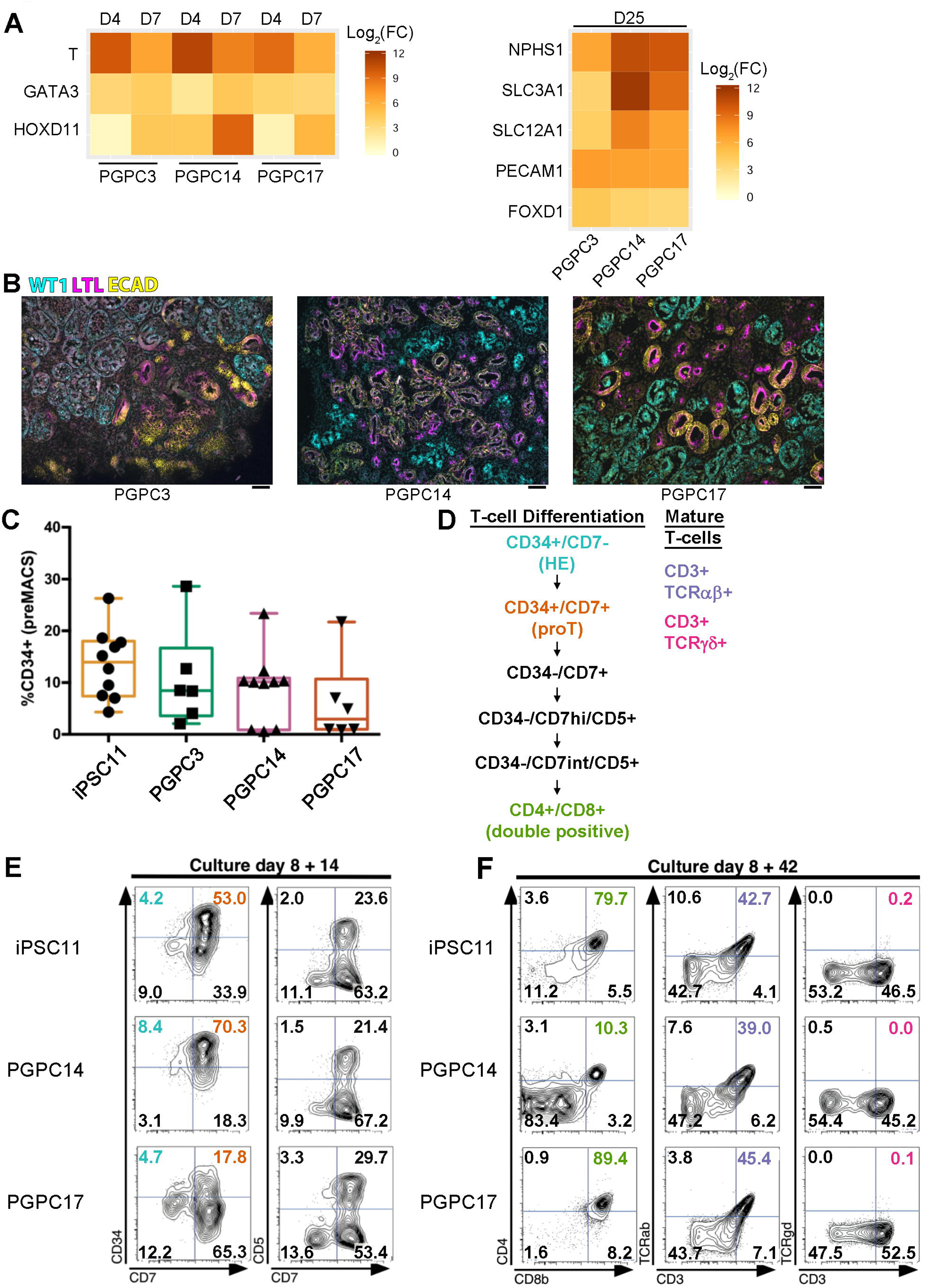
Generation of kidney organoids and T-lymphocytes. (A) Heatmaps indicating log_2_ fold change of marker gene expression normalized to iPSCs (left) or D7 (right) [biological replicates (PGPC3/17 = 1, PGPC14 ≥ 2); technical replicates ≥ 3]. (B) Representative images of immunohistochemistry of kidney organoid sections labeled with anti-WT1, anti-LTL, and anti-ECAD antibodies (biological replicates ≥ 3 for each line). Scale bar represents 100 *µ*m. (C) Average differentiation efficiency of iPSC lines after MACS sorting for CD34 (biological replicates = 4; SEM). (D) T-lymphocyte marker expression during T-cell maturation from CD34+ HE cells. (E) D8+14 cells were labeled with anti-CD34, anti-CD7 and anti-CD5 antibodies and analyzed by flow cytometry (biological replicates = 3). (F) D8+42 cells were labeled with anti-CD4, anti-CD8b, anti-CD3, anti-T-cell receptor alpha/beta, and anti-T-cell receptor gamma/delta antibodies then analyzed by flow cytometry (biological replicates = 3).

To evaluate the potential to generate hematopoietic stem/progenitor cells (HSPCs) and T-cells, we compared the PGPC lines to iPSC11 (Alstem Cell Advancements) in an embryoid body differentiation protocol described by Kennedy et al., 2012 with feeder-free adaptation (Fig. S2B). All three PGPC lines gave rise to CD34+ hemogenic endothelial (HE) cells with a similar efficiency as iPSC11 cells (Fig. 4C), which were magnetic-activated cell sorted at D8 (Fig. S2C). PGPC14 and 17 were most enriched for CD34+ cells, which were then cocultured on OP9-DL4 cells to induce a T cell differentiation program. Next, multi-colour flow cytometry was used to simultaneously measure different cell populations at D8+14 and D8+42 to assess the ability of HE-derived HSPCs to differentiate to T-lymphocytes, a hallmark of definitive hematopoietic potential (Fig. 4D). At D8+14 (Fig. 4E), early T-lineage progenitor cells (proT cells marked as CD34+ CD7+) could be observed transitioning to more differentiated early T-lineage cells (CD34-CD7+), with a subpopulation co-expressing a pan-T cell marker (CD7+ CD5+). PGPC14 exhibited a prolonged proT stage (70%), whilst PGPC17 showed a more rapid transition (17%) compared to iPSC11 (53%). Simultaneous assessment of T-cell markers on culture day 8+42 detected the presence of double positive (DP) CD4+ CD8+ T-lineage cells, expressing T-cell receptors (TCR/CD3+). At this time point, the PGPC lines showed similar propensity as iPSC11 to generate TCR alpha/beta (39-45%) cells, but only PGPC17 produced rare TCR gamma/delta bearing (0.1%) cells. We conclude that PGPC lines differentiate into HE cells that develop into into T-cells but with different differentiation kinetics that may require line-specific protocol optimization.

### Utility of the resource - Sensory neuron protocol optimization and subtype identification

PGPC17_11 was selected to optimize differentiation into peripheral sensory neurons (PSNs) using a small molecule inhibitor protocol adapted from (Chambers et al., 2012) (Fig. 5A). We found improved growth and survival with the addition of NGF at D2 and using N2 media with the following composition: 47.5% DMEM/F12, 47.5% neurobasal, 2% B-27 supplement, 1% N2 supplement, 1% glutamax and 1% Pen/Strep (full methods described in supplemental experimental methods). Whole cell patch clamp recordings were used to assess excitability. At two weeks post-induction, all PSNs responded to sustained current injection with transient spiking but, by four weeks, half the neurons had switched to repetitive spiking (Fig. 5B). The action potential waveform also experienced significant changes, which included an increase in amplitude (Fig. 5C) and a decrease in width (Fig. 5D) amongst both transient and repetitive spiking 4-week old neurons compared with 2-week old neurons. Amongst 4-week old neurons, repetitive spiking neurons had a significantly lower rheobase (current threshold) than transient spiking neurons (Fig. 5E). Additional membrane properties are described in Fig. S3A-F

**Figure 5.**
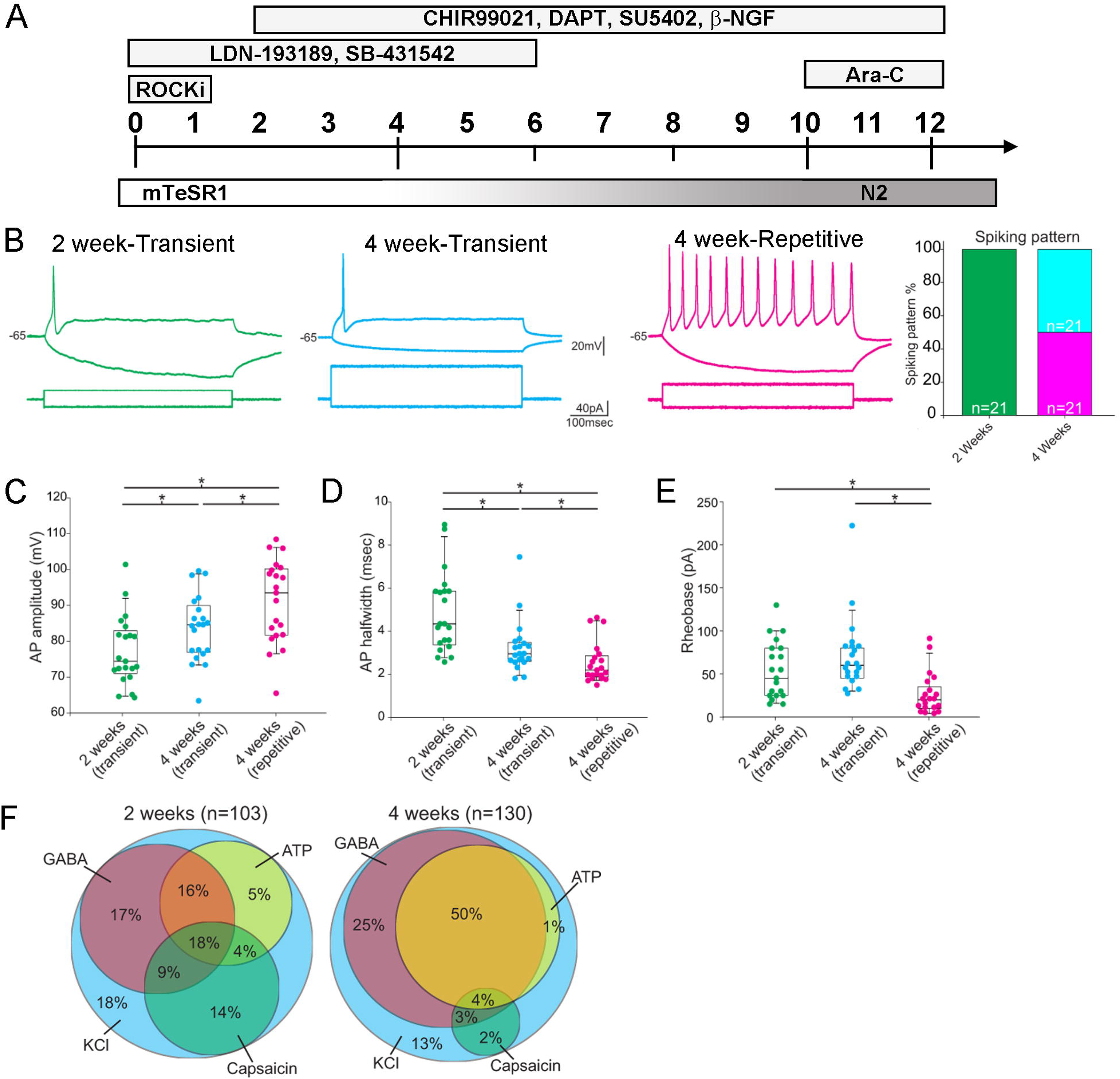
PGPC17-derived sensory neurons are predominately non-peptidergic. (A) Sensory neuron differentiation scheme. The LDN and SB drug combination was applied between D0 through D5 with the CHIR, DAPT, SU and NGF combination starting on D2 through D11. Starting on D4, N2 media was added in increasing 25% increments replacing mTeSR1. Dividing cells were eliminated using Ara-C on D10. N2 media was changed twice weekly thereafter. (B) Representative spiking patterns to sustained somatic current injection. At 2 weeks, all cells spike transiently (biological replicates = 3; technical replicates ≥ 5) whereas at 4 weeks, there is a significant increase in the proportion of repetitively spiking neurons (biological replicates = 4; technical replicates ≥ 2) compared to transiently spiking neurons (biological replicates = 4; technical replicates ≥ 2) (p<0.0001, chi square test). The action potential waveform experienced a significant increase in amplitude (C) and a significant decrease in width (D) between 2- and 4-weeks post-induction. Rheobase was significantly lower in repetitive spiking neurons than in transient spiking neurons at 2- or 4-week post-induction (E). * shows p < 0.05 based on Mann-Whitney U tests. (F) Ca^2+^ revealed a significant decrease in the proportion of neurons responsive to capsaicin between 2 and 4 weeks (<0.00001, chi square test) whereas the proportion of neurons responsive to GABA (p=0.0001) or ATP (p=0.042) significantly increased.

To further characterize phenotype, we imaged the Ca^2+^ responses evoked by brief application of various agonists. Neurons exhibiting a robust Ca^2+^ response to KCl application were considered healthy and their responses to capsaicin, GABA, and ATP were tested (Fig. 5F). At two weeks post-induction, 44.6% of neurons responded to the TrpV1 agonist capsaicin, but that number fell to 10% by week four (p < 0.00001). TrpV1 is a marker of peptidergic nociceptors, but is broadly expressed among immature PSNs and is developmentally downregulated (Cavanaugh et al., 2011). Our data suggest that iPSC-derived PSNs follow a similar developmental program. Low TrpV1 expression at four weeks suggests that repetitive spiking PSNs represent predominantly non-peptidergic nociceptors (Zeisel et al., 2018), whereas the transient spiking neurons are most likely mechanoreceptors. The proportion of neurons responsive to GABA increased over time (p = 0.0001), as did the proportion responsive to the purinergic receptor agonist ATP (p = 0.042). These results demonstrate that PGPC17 was successfully differentiated into active neurons with a non-peptidergic nociceptor or mechanoreceptor phenotype.

### Utility of the resource - WGS analysis

To identify iPSC-specific variants we obtained whole genome sequences of each PGPC line to compare to their respective donor blood sequences (Table 1). On average, we identified 1,462 novel nucleotide variants [range: 1,169-1,981] and 0.3 novel copy number variants [range: 0-1] per clone. Twenty variants [range: 18-23] affected exonic regions: 13 were non-synonymous [range: 12-14] and 1.3 loss-of-function [range: 1-2]. One likely pathogenic *TP53* variant (Arg158His) was called in PGPC3_75 at low allelic fraction (4/22 reads), compatible with mosaicism (Table S2). We did not identify any other known pathogenic sequence variants in reprogrammed cell lines. Three loss-of-function variants, although not associated with human disease, were in genes with high haploinsufficiency scores and known function in embryonic development (PGPC14_26: *TRIM71* and *FRMD4A*, PGPC17_11: *ROBO2*, Table S3). For PGPC14_26, we also identified an intronic 16-kb-deletion of uncertain significance in *IL1RAPL1*, a gene associated with impaired synaptogenesis and neurodevelopmental deficits. Annotation of the PGP donor and cell-line derivative sequences will be important as variant databases mature (Costain et al., 2018).

### Utility of the resource-CRISPR/Cas9 gene editing and phenotyping

To edit a gene for cardiac phenotyping, we targeted a region of *MYBPC3* where frameshifts are associated with hypertrophic cardiomyopathy by using guide (g)RNAs for CRISPR-Cas9 directed non-homologous end-joining. We nucleofected PGPC17_11 iPSCs with a pSpCas9(BB)-2A-Puro vector containing gRNA sequences targeting *MYPBC3* (Fig 6A). Transfected cells were selected with puromycin treatment and resistant colonies were isolated and expanded. A karyotypically normal sub-clone bearing an apparent homozygous frameshift mutation was identified in the *MYBPC3*_KO line by Sanger sequencing (Fig. 6A/B). To characterize genetic changes, we performed WGS. On-target compound heterozygote *MYBPC3* frameshifts were shown to be an 8-bp insertion at chr11:47,359,282insGTGCAGGA, and a large >260 bp insertion at the same position in the other allele. This insertion did not map to the human genome and was not detected using our PCR-based sequencing due to the size of the insertion. To characterize potential off-target effects in the *MYBPC3*_KO cells, we first used benchling.com’s prediction tool to identify the top 49 off-target sites. We searched 100 base-pairs up and downstream of each predicted site and found zero novel variants within these regions. When we looked for overall novel genomic variation, 917 new single nucleotide variants and one intergenic 32 kb deletion (chr18:12137685-12169689) were found (Table 1). None of these variants were likely pathogenic, similar to our other reported gene edited lines (Deneault et al., 2018, 2019; Zaslavsky et al., 2019).

**Figure 6.**
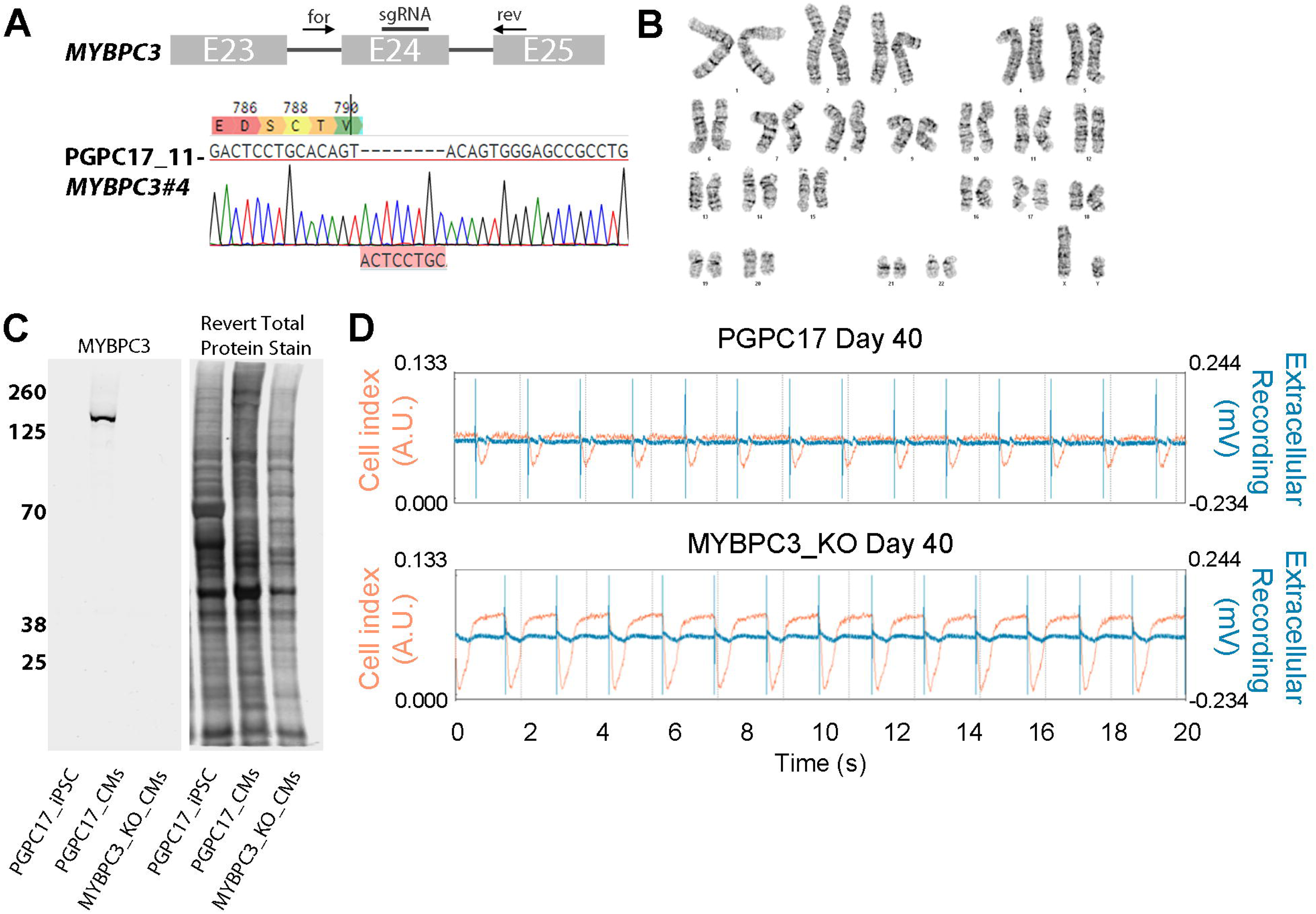
Derivation of *MYBPC3*-knockout iPSC that display a cardiomyopathy phenotype. (A) Exon 24 of *MYBPC3* was targeted by CRISPR/Cas9 and Sanger sequencing of one clone PGPC17_11-*MYBPC3*#4 (hereby identified as MYBPC3_KO) identified an out of frame insertion resulting in an early stop codon. (B) Normal karyotype was confirmed before differentiation and characterization. (C) Western blot probing for MYBPC3 using iPSC lysate as a negative control and parental CM lysate as a positive control. No full length or truncated forms of MYBPC3 were detected on the blot using near infrared detection. Revert total protein stain was used to show similar amounts of protein were added to each lane. (D) Representative xCELLigence traces of PGPC17 and MYBPC3-KO CMs showing beat amplitude (cell index) and extracellular voltage recordings over a 20 second sweep at D40 (biological replicates = 2; technical replicates ≥ 8). MYBPC3-KO CMs showed a higher beat amplitude compared to isogenic control CMs suggestive of hypercontractility as seen in hypertrophic cardiomyopathy.

To examine the consequences of the frameshifts on MYBPC3 protein, we generated CMs as described in Figure 3 and collected protein lysates from PGPC17 parental and *MYBPC3_KO* iPSC-CMs. Western blots were unable to detect MYBPC3 protein in the KO clone (Fig. 6C). We matured CMs until D36-44 to look for phenotypic evidence of hypertrophic cardiomyopathy as predicted by loss of MYBPC3. Indeed, xCELLigence assays detected increased BAmp in the MYBPC3_KO-CMs compared to the parental line at D42 (0.08 AU and 0.04 AU respectively) while having similar beat rates (41 bpm to 36 bpm respectively). Recently, Cohn et al., 2019 generated a frameshift in *MYBPC3* using cells from the American PGP and observed similar phenotypes. Our findings demonstrate the utility of PGPC17_11 for gene editing to produce isogenic cell lines for cardiac phenotyping.

### WGS analysis of publicly available HipSci lines

Since we found that all our iPSC lines have pre-existing and/or novel variants of potential concern when considering experiments for different lineages, we analyzed downloaded genome sequencing data of five publicly available HipSci lines suggested as healthy controls (HPSI0114i.kolf_2, HPSI0214i.kucg_2, HPSI0214i.wibj_2, HPSI0314i.hoik_1 and HPSI0314i.sojd_3). Across all five samples, 89-96% of the genome was covered at least 20× (quality metrics in Table S4). We interpreted likely pathogenic variants, loss-of-function constraint gene variants and variants of unknown significance (VUS) as previously described (Reuter et al., 2018) (Supp. Excel files).

Two likely pathogenic variants were found in kolf_2 and one in sojd_3 that were predicted to have clinical relevance if identified in humans and could also affect experimental assays. kolf_2 had a substitution of two adjacent nucleotides, disrupting exon-intron boundaries of one *COL3A1* allele. One variant was within canonical splice site c.3526-1G>A, and likely to cause out-of-frame exon skipping. If splicing was preserved, the second nucleotide change would result in a likely pathogenic missense alteration p.(Gly1176Ser). COL3A1 haploinsufficiency is associated with dysfunctional connective tissue, such as in the vascular system, skin, intestine, lung, and uterus, and causes vascular type (IV) Ehlers-Danlos syndrome. The same kolf_2 line also harboured a heterozygous 19-basepair deletion p.(Pro197Hisfs*12) in *ARID2*. The variant was likely pathogenic for Coffin-Siris syndrome, a neurodevelopmental disorder with variable skeletal and organ manifestations. Finally, sojd_3 harboured a likely pathogenic heterozygous nonsense variant p.(Gln348*) in *BCOR*. This X-linked gene encodes a transcriptional corepressor with important functions in early embryonic development of various tissues (Wamstad et al., 2008). Females with heterozygous *BCOR* defects may exhibit oculofaciocardiodental syndrome. None of these likely pathogenic variants had been previously reported, and we cannot determine if they were present in the donor genomes, or arose during reprogramming, and could therefore be mosaic. We also identified several loss-of-function variants of uncertain significance in constrained genes in hoik_1 and kucg_2, mostly with known functions in early development (as in *PTK2, ZNF398, UBE3C, CDC37, TNS3*; Table S3) and many VUS (Supp Excel files). We did not identify pathogenic or likely pathogenic variants in wibj_2 which suggests it is the variant-preferred line amongst this subset.

## DISCUSSION

Here we generated a resource of iPSC control lines for use in disease modeling studies. These cells have the benefit of both annotated genomic variants and demonstrated multilineage directed differentiation into functional cortical neurons, cardiomyocytes and hepatocytes. Pilot users showed that the lines can be used to generate kidney organoids, or T-lymphocytes to identify specific subtypes of active sensory neurons, and for gene editing which revealed a preliminary phenotype in isogenic *MYBPC3* KO cardiomyocytes similar to another isogenic pair (Cohn et al., 2019).

Apart from their versatility, the main advantage of these blood-derived footprint-free lines is the clinical annotation of potentially disease-associated variants that may impact cellular phenotypes. Variant analysis in the PGPC participants’ blood had revealed heterozygous variants of unknown significance in all individuals (Reuter et al., 2018). This observation suggests that it may not be possible to isolate universal control lines and reinforces the importance for WGS in characterizing control lines, especially as clinical annotation gains precision with ongoing variant discoveries. WGS has the advantage of allowing detection of coding variants, CNVs and non-coding variants although the latter have not yet been fully explored in these lines. Knowledge of the donors’ genomes allowed predictions on how to prioritize control lines for use as tissue specific controls. For example, as PGPC3 and PGPC14 had variants that could predispose to altered cardiac channel function, PGPC17 was deemed to be the preferred line for the study of cardiac disease. PGPC3 however was preferred for neurological disorders.

WGS has previously determined that iPSC lines have variants that differ from those in the donor. Our WGS data reveals that the reprogrammed lines have more than a thousand new SNVs each, whereas only 1 new CNV (15.6 kb) was detected likely due to prior selection for normal karyotype. Most variants were of uncertain significance, with three new variants of potential concern in PGPC14_26 (Table S3). Genome sequencing of the *MYBPC3* KO line showed more than 900 additional SNVs compared to the unedited iPSC line. None of the new variants were near potential gRNA cut sites, suggesting they were not off-target and were indeed de novo mutations. These analyses highlight that iPSC lines harbor variants of potential concern that are not found in the donor blood. Moreover, our annotation of 5 healthy control lines from the HipSci consortium discovered likely pathogenic variants in 2 lines and additional loss-of-function variants in constrained genes in 2 other lines, leaving only wibj_2 as a preferred healthy control line. We propose that clinical annotation of WGS data is an important quality control measure of iPSC lines, and that its expanded use will identify additional variant-preferred lines to use as healthy controls for disease modeling.

Disease modeling has generally used 2-3 lines from each individual to account for variability in reprogramming. To account for 1000-2000 novel variants in each line compared to the parental genome, this study provides another rationale for studying multiple lines from each individual. With this in mind, we generated a resource of 4-5 iPSC lines each from two males and two females, all with standard pluripotency characterization available. We additionally performed multilineage directed differentiation on a single line from three individuals, assuming that single lines from 3-4 individuals can account for inter-individual variability. One highly characterized line is therefore available from three PGPC participants, and preferred lines are likely to be of high utility for gene editing studies that compare the phenotype of isogenic cells. Ultimately, users of the resource will select one or more lines from each PGPC participant depending on their research strategy.

## EXPERIMENTAL METHODS

Reprogramming of PGPC iPSCs was performed under the approval of the Canadian Institutes of Health Research Stem Cell Oversight Committee, and the Research Ethics Board of The Hospital for Sick Children, Toronto. Blood cells were reprogrammed with Sendai virus to deliver reprogramming factors, and iPSCs were maintained in feeder-free conditions with mTeSR1 (STEMCELL Technologies); see Supplemental Information. WGS was performed on Illumina HighSeq X and analyzed as previously described (Reuter, et al. 2018). A vector-based CRISPR/Cas9 approach was used to mutagenize *MYBPC3*, further described in Supplemental Information. Detailed descriptions of differentiations, characterizations and functional assays are found in Supplemental Information. Over expression of NGN2 induced iPSCs to differentiate to glutamatergic neurons. Extracellular electrophysiology recordings were collected with an Axion Maestro MEA reader (Axion Biosystems) multi-electrode array as described in the Supplemental Information. CMs were differentiated using STEMdiff Cardiomyocyte differentiation kits (STEMCELL Technologies). CM Calcium imaging was captured by loading cells with Fluo-4 dye and taking images at 4 Hz for 30 seconds. Contractile and electrical activity was recorded with an xCELLigence RTCA CardioECR (ACEA Biosciences). HLCs were generated as previously described (Ogawa et al., 2015). CYP3A4/7 was measured using p450-Glo assay kit (Promega) as per manufacture’s protocol. Kidney organoids were generated as previously described (Takasato et al., 2015). T-cells were generated as previously described (Kennedy et al., 2012; La Motte-Mohs et al., 2005). Sensory neurons were generated using a modified method described in Chambers, et al. 2012. Whole cell electrophysiology recordings where made at room temperature with an Axopatch 200B (Molecular Devices) from borosilicate patch electrodes. Ca^2+^ imaging was performed on neurons incubated in Ca^2+^ green-1 AM dye (ThermoFisher Scientific) at room temperature. Images were acquired at 25 Hz using a NeuroCCD-SM256 imaging system (RedShirt Imaging). WGS datasets are available from EGA (EGAS00001003684) and RNAseq datasets are available from GEO (<u>GSE132012</u>). iPSC lines are available upon request.

## AUTHOR CONTRIBUTIONS

M.R.H., M.S.R., S.W.S. and J.E. designed the research project. M.R.H., W.W., N.T., J.L., S.S., J.M., L.S.L., P.M.B., A.P., A.R. and G.M. contributed to cell maintenance, characterization, and differentiation. M.R.H., N.T. and W.W. contributed to the CRISPR experiments. M.S.R. performed clinical annotation and off-target analyses. C.K., D.C.R., J.L.H., P.P., R.S.M., M.R., E.C.M.P., and M.J.S. provided technical help. M.R.H., M.S.R., N.T., J.L., J.M., L.S.L., P.M.B., J.C.Z-P., M.K.A., S.A.P., N.D.R., B.M.K., S.M., S.W.S. and J.E. wrote the manuscript with comments from all co-authors. M.R.H. and J.E. supervised the project. Specific contributions of the co-corresponding authors: J.C.Z-P. and M.K.A. labs differentiated iPSCs to T-cells and conducted flow cytometry; S.A.P. lab conducted patch-clamp electrophysiology and Ca^2+^ imaging experiments on sensory neurons; N.D.R. lab differentiated iPSCs to kidney organoids and performed expression analysis and imaging; B.M.K. lab differentiated iPSCs to HLCs and performed imaging, expression and enzymatic analysis; S.M. lab performed CM ICC and Ca^2+^ imaging and xCELLigence analysis, S.W.S. lab performed WGS analysis and annotation; J.E. lab generated iPSCs and differentiated iPSCs to sensory neurons and CMs for other groups and cortical neurons for imaging and multi-electrode arrays.

## ACKNOWLEDGEMENTS

The research was supported by grants from the McLaughlin Centre (MC-2014-06) and Ontario Brain Institute Province of Ontario Neurodevelopmental Disorders Network (J.E., S.W.S.; IDS-11-02), GlaxoSmithKline - Canadian Institute of Health Research (CIHR) Chair in Genome Sciences (S.W.S.), Ted Rogers Centre for Heart Research Strategic Innovation grant (J.E. and S.M.), Heart and Stroke Foundation Chair (S.M.), CIHR Team Grant (B.M.K.; THC-135232), Tier I Canada Research Chair and CIHR Foundation Grant (N.D.R.; SOP-155609), CIHR Chronic Pain Network-SPOR (S.A.P. and J.E.; 2017-007), Medicine by Design New Ideas grants (J.E., M.K.A.; MBDNICL-2017-03 and J.C.Z-P., M.K.A.; C1TPA-2016-20), NSERC RGPIN grant (M.K.A.; 05333-14), and Fellowship support from SickKids RestraComp award (J.M and S.S.). We thank B. Thiruvahindrapuram, T. Nalpathamkalam, W.W. Sung, Z. Wang, and G. Kaur for bioinformatic support; the Centre for Applied Genomics, the SickKids-UHN Flow and Mass Cytometry Facility and the SickKids Imaging Facility for technical support. The NGN2/rtTA lentiviral constructs were gifts from T.C. Südhof, and we thank N.N. Kasri and K. Linda for technical advice. We thank the Personal Genome Project Canada, Cheryl Cytrynbaum, Ny Hoang and Barbara Kellem for genomic data and collecting samples for reprogramming, and the PGPC blood donors for volunteering to participate in this research.

