## Supplemental_data for "Control iPSC lines with clinically annotated genetic variants for versatile multi-lineage differentiation"

SUPPLEMENTAL EXPERIMENTAL PROCEDURES

*Reprogramming and cell culture*	Reprogramming of iPSCs was performed under the approval of the Canadian Institutes of Health Research Stem Cell Oversight Committee, and the Research Ethics Board of The Hospital for Sick Children (REB#1000050639). CD34+ cells were isolated from blood samples collected from each of the four PGPC donors using CD34+ Reprogramming Kit (STEMCELL Technologies) according to the manufacture’s protocol. Cultured CD34+ cells were tested for mycoplasma by PCR (see Table S7 for primer pairs) during routine early passaging then reprogrammed using CytoTune-iPS 2.0 Sendair Reprogramming Kit (ThermoFisher) with modifications post transduction. Transduced cells were plated in ReproTeSR (STEMCELL Technologies) and isolated colonies were directly transferred to mTeSR1 (STEMCELL Technologies).

*iPSC maintenance*	All iPSC lines were maintained on Matrigel (Corning) coated dishes using mTeSR1. Daily media changes were performed except for the day following passaging. ReLeSR (STEMCELL Technologies) was used to lift clumps of iPSCs for routine passaging. Accutase (InnovativeCellTechnologies) and 10 µM Rho-associated kinase (ROCK) inhibitor (Y-27632; STEMCELL Technologies) were used for single-cell dissociation purposes unless otherwise specified. Mycoplasma was routinely tested as cells were maintained.

*PCR for Sendai virus*	Total RNA was isolated using TRIzol (Invitrogen) according to the manufacture’s protocol. 1 µg isolated RNA was used to generate cDNA with SuperScript VILO cDNA Synthesis kit (ThermoFisher) according to the manufacture’s protocol. 10 µl of the reaction was amplified with AccuPrime *Taq* DNA polymerase with Sendai virus primers Table S7. Resulting DNA was run on a 2% agarose gel, stained with ethidium bromide and imaged. Passage (P)2 iPSC cDNA samples were used as a positive control for Sendai virus.

*Karyotyping*	Karyotyping and standard G-banding chromosome analysis with 400-banding resolution was performed at The Centre for Applied Genomics (TCAG).

*AR assay*	400 ng genomic DNA was digested with HpaII and HhaI simultaneously for 5 hours. 2 µl digest was used in a PCR with Platinum™ *Taq* DNA Polymerase High Fidelity (Invitrogen) using *AR* gene primers (Table S7). 15 µl PCR was sent to TCAG for Genetic Analysis electrophoresis and resulting traces were analyzed using Peak Scanner software. Male samples were used to confirm complete digestion.

*RNA sequencing and Pluritest*	RNA was collected from single wells of 6-well plates using RNA PureLink RNA mini kit (Life Technologies) as per supplied protocol. RNA samples were sent to TCAG where RNA quality was checked by Bioanalyzer then sequenced as paired end 2x125 bases on an Illumina HiSeq 2500 at a depth of 20M reads. Resulting compressed FASTQ files were uploaded to www.pluritest.org for analysis.

*Immnocytochemistry*	IPSCs were plated on 24-well dishes for epifluorescence microscopy while differentiated cells were seeded on µ-Plate 24-well Black plates (IBIDI) for confocal microscopy. IPSCs, iPSC-derived cardiomyocytes and iPSC-derived hepatocytes were fixed with 4% paraformaldehyde (PFA), while iPSC-derived neurons were fixed with PKS (4% PFA in Krebs-Sucrose buffer, 50 mM KCl, 1.2 mM CaCl_2_, 1.3 mM MgCl_2_, 20 mm HEPES pH 7.4, 12 mM NaH_2_PO_4_, 400 mM sucrose, 145 mM NaCl, 10 mM glucose in water). Three washes with phosphate buffered saline (PBS) was performed followed by permeabilization with 0.1% Triton X-100 incubation for 10 minutes. Cells were blocked with blocking solution for one hour. Primary antibody (for appropriate dilutions see Table. S8) was prepared in blocking solution and incubated with the cells at 4^o^C overnight. Three washes with PBS were done before incubating with fluorescence-tagged secondary antibodies (Table S8) for one hour in the dark at room temperature. Cells were washed twice with PBS then incubated with 1:2000 4′,6-diamidino-2-phenylindole (DAPI) for 5 minutes to counterstain nuclei followed with a final wash of PBS.

*Imaging*	IPSCs and sensory neurons were imaged on a Leica DM14000B epifluorescence microscope using a DFC 7000T camera and LAS X software (Leica). Cortical neurons were imaged with an Olympus IX81 spinning disk confocal using a Hamamatsu C9100-13 EM-CCD camera and Volocity software (Perkin Elmer). Cardiomyoctyes were imaged with a Zeiss AxioVert 200M microscope using a Hamamatsu C9100-13 EM-CCD camera and Volocity software. Hepatocytes and kidney organoid sections were imaged with a Zeiss AxioVert 200M microscope, using an AxioCam HRm camera, and AxioVision (V4.9.2 SP3). ImageJ (V1.52i) and Photoshop (V12.0) were used for image analysis and figure preparation.

*Whole genome sequencing (WGS) and clinical annotation*	Genomic DNA was isolated using Quick-DNA miniprep kit (Zymo Research) according to manufacturer’s protocol. 1 µg gDNA was submitted to TCAG for genomic library preparation and whole genome sequencing on an Illumina HighSeq X system as described in Reuter et al., 2018. For reprogrammed PGPC lines, we performed genome sequencing as previously described (Reuter et al., 2018). We used Illumina provided software bcl2fastq (v2.20) to convert the per-cycle BCL basecall files to standard sequencing output in FASTQ format. FastQC was used to assess the quality of the experiment, and FastQ Screen to check the composition of the library. Reads were aligned to the reference human genome (GRCh37) using BWA mem (v0.7.12) (Li and Durbin, 2009). Duplicate reads were marked using Picard Tools (v2.5.0). Indel realignment, base quality score recalibration and germline variant detection using HaplotypeCaller were performed using GATK 3.7 following the best practices recommendation (Van der Auwera et al., 2013; DePristo et al., 2011). Resulting variant calls were annotated using a custom pipeline developed at TCAG (Yuen et al., 2015), based on ANNOVAR (Wang et al., 2010). ERDS (v1.1) (Zhu et al., 2012) and CNVnator (v0.3.2) (Abyzov et al., 2011) were used to call CNVs, and a custom annotation and prioritization pipeline was used to define rare CNVs (Reuter et al., 2018). We used Mutect2 (v3.7-0, default parameters) to detect somatic variants in the reprogrammed cell lines that were not found in blood (Cibulskis et al., 2013). Variants were annotated using an ANNOVAR based pipeline (Wang et al., 2010).
We downloaded genome sequencing data (fastq files) of five publicly available iPS cell lines: HPSI0114i.kolf_2, HPSI0214i-wibj_2, HPSI0214i-kucg_2, HPSI03145i-hoik_1, HPSI0314i-sojd_3 (http://www.hipsci.org/lines/#/lines, https://www.ebi.ac.uk/ena). We performed data analysis, variant prioritization and interpretation as described previously (Reuter et al., 2018).

*CRISPR-Cas9 gene editing*
pSpCas9(BB)-2A-Puro (PX459) V2.0 (Addgene #62988) was cloned with oligos designed to target *MYBPC3* exon 24 using benchling.com CRISPR prediction tool (Benchling, 2019). Indels were introduced into iPSCs by nucleofecting 2x10^6^ cells with 5 µg plasmid using Nucleofector 2b (Lonza) with program A-23 (~24% transfection efficiency). Transfected cells were split to multiple wells of a 6-well plate. On D2, puromycin (0.5 ug/ml) was added to daily mTeSR1 changes until D5. Surviving single colonies were grown to D12-18 to ensure they were large enough to isolate and transfer to 24-well plates. Isolated clones were passaged and genomic DNA (gDNA) was harvested from residual cells after the first 6-well passage with Quick DNA Miniprep kit. gDNA was genotyped by submitting PCR products (Table S7) for Sanger sequencing by TCAG.
Subsequent editing of other genes was performed with 8x10^5^ cells and 1.5 µg plasmid in 100 µl scale Neon Transfection System (ThermoFisher) with one pulse at 1500 millivolts for 30 milliseconds (~40-70% transfection efficiency). Nucleofected cells were plated in 2 wells of a 6-well plate with mTeSR1 media supplemented with CloneR (STEMCELL Technologies) to enhance cell survival.

*NGN2-Lentivirus preparation*	Two T-75 flasks were seeded with 7.5x10^6^ HEK293-T cells and grown in DMEM (Gibco) supplements with 10% fetal bovine serum (Gibco) and antibiotics penicillin and streptomycin. Cells were transfected 24 hours post seeding with Lipofectamine 2000 (Thermofisher) with 10 µg gag-pol, 10 µg rev, 5 µg VSV-G and 15 µg *NGN2* or rtTA plasmids (Hotta et al., 2009). Media was removed 18 hours later and increased to 25 ml per dish. The following day, supernatant containing viral particles were collected, filtered and concentrated by 91,000 g centrifugation for 2 h at 4°C. The supernatant was discarded and 50 µl HBSS was added to the pellet and left overnight at 4^o^C. The following day, aliquots of 10 µl were frozen at -80^o^C.

*Differentiation and maintenance of excitatory cortical neurons*	Excitatory cortical neurons were generated as previously described by Deneault et al. 2018. In brief: 5x10^5^ iPSCs/well were seeded in Matrigel-coated 6-well plate in 2 ml of mTeSR1 supplemented with 10 μM Y-27632. The following morning media in each well was replaced with fresh mTeSR1 plus 10 μM Y-27632, 0.8 μg/ml polybrene (Sigma), and minimal amounts of *NGN2* and *rtTA* lentiviruses necessary to generate >90% GFP+ cells upon doxycycline induction. Viral volumes were determined for each virus batch by titration and flow cytometry measurement of GFP positivity. The day after transduction, virus-containing media were replaced with fresh mTeSR1, and cells were expanded to 80-90% confluency. *NGN2*-iPSCs were expanded once to freeze aliquots of infected cells. *NGN2*-iPSCs reliably generated neurons up to passage seven.
*NGN2*-iPSCs were dissociated using Accutase and seeded in Matrigel-coated 6-well plates at a density of 5x10^5^ cells per well in 2 ml of mTeSR1 supplemented with 10 μM Y-27632 (D0). D1, media in each well was changed for 2 ml of CM1 [DMEM-F12 (Gibco), 1x N2 (Gibco), 1x NEAA (Gibco), 1x pen/strep (Gibco), 1 µg/ml laminin (Sigma), 10 ng/µl BDNF (Peprotech) and 10 ng/µl GDNF (Peprotech)] supplemented with 2 µg/mL doxycycline hyclate (Sigma) and 10 μM Y-27632. D2, media was replaced with 2 ml of CM1 supplemented with 2 µg/ml doxycycline hyclate and 2 µg/ml puromycin (Sigma). D3 media was changed to CM2 [Neurobasal media (Gibco), 1x B27 (Gibco), 1x Glutamax (Gibco), 1x pen/strep, 10 ng/µl BDNF and 10 ng/µl GDNF] supplemented with 2 µg/ml puromycin and 1 µg/ml laminin. The same media change was repeated at D4 without puromycin. D6, media was replaced with CM2 supplemented with 2 µg/ml doxycycline hyclate, 10 µM Ara-C (Sigma), 10 ng/µl BDNF, 10 ng/µl GDNF, and 1 µg/ml laminin. D8 post-*NGN2*-induced neurons were dissociated with Accutase filtered through a 70 µm filter and seeded for subsequent experiments, as described below.

*Seeding neurons for imaging*	Two days before seeding neurons, 24-well black µ-Plates were coated overnight with poly-L-ornithine followed by an overnight coating with 40 µg/ml laminin. 3.5x10^5^ neurons were seeded in each well in CM2 media and 24 hours later 7x10^4^ mouse astrocytes/well were seeded on top of neurons. Astrocytes were prepared from postnatal day 1 CD-1 mice as described (Kim and Magrane, 2011). Before use, astrocytes were checked for mycoplasma contamination. All animal work was approved by The Hospital for Sick Children Animal Care Committee and complies with the guidelines established by The Canadian Council of Animal Care. CM2 media was replaced twice weekly. 48 hours prior to fixation and staining neuron we sparsely transfected each well with 1 µg pmax-FP-Green-N plasmid carried by Lipofectamine 2000 in Opti-MEM. 16-18 hours post transfection media was removed and replaced with CM2 media. Neurons were prepared and imaged as described above.

*Tracing neurons*	Blinded images were analyzed using Imaris (Bitplane). Somas and dendrites were manually traced using the GFP channel. Axonal tracing was prevented by using the MAP2 channel to identify dendrites. 40 neurons were measured from each cell line (20 technical replicates; 2 separate differentiations).

*Seeding neurons for MEA recording*	48-well opaque- or clear-bottom MEA plates (Axion Biosystems) were coated with filter sterilized 0.1% PEI solution in borate buffer pH 8.4 for 1 hour at room temperature, washed four times with water, and dried overnight. 1.2x10^5^ D8 neurons were seeded per well as 40 µl droplets in CM2 (BrainPhys supplemented with BDNF, GDNF) with 400 µg/µl laminin and allowed to sit for 1 hour at 37^o^C before adding 300 μl CM2 supplemented with 40 ug/ul laminin. 24 hours later, 1x10^4^ mouse astrocytes/well were seeded on top of neurons in CM2. Media was changed twice a week on the same days of the week. Once a week from week four to seven, electrical activity of the MEA plates was recorded using the Axion Maestro MEA reader (Axion Biosystems). Heater control was set to 37°C. Each plate was incubated for five minutes on the warmed reader, then real-time spontaneous neural activity was recorded for five minutes using AxIS 2.0 software (Axion Biosystems). A bandpass filter from 200 Hz to 3 kHz was applied. Spikes were detected using a threshold of 6 times the standard deviation of noise signal on electrodes. Offline advanced metrics were re-recorded and analyzed using Axion Biosystems Neural Metric Tool. Electrodes were considered active if at least 5 spikes were detected per minute. Bursts were considered as groups of at least 5 spikes, each separated by an inter-spike interval (ISI) of no more than 100 ms. Network bursts were identified by a minimum of 10 spikes with a maximum ISI of 100 milliseconds covered by at least 25% of electrodes in each well. No non-active well was excluded in the analysis. After the last reading at week seven, selected wells were treated with a synaptic antagonist to AMPA receptor: 6-cyano-7-nitroquinoxaline-2,3-dion (CNQX; Sigma) at 60 μM. Plates were recorded 15 minutes after addition of the antagonists. Culture media was replaced with CM2 and after a one-hour recovery period a final recording was taken.

*Statistical analysis of neurons*	All statistical tests were conducted in RStudio (V1.1.456). Comparisons between soma area and dendritic length were conducted using Kruskal-Wallis tests after determining non-normal distribution of the data by qqplots and a Shapiro test. Comparisons between cell lines at each distance of Sholl analysis was conducted using Dunn’s test when finding statistical significance by a Kruskal-Wallis test.

*Differentiation and maintenance of CMs*	Cardiomyocytes were generated using STEMdiff Cardiomyocyte Differentiation Kit (STEMCELL Technologies). iPSCs were dissociated with Accutase and seeded at 8x10^6^ cells per Matrigel-coated well of a 12-well dish in mTesR1 supplemented with 10 µM Y-27632. 24 hours later media was replaced with mTesR1. 48-72 hours post-seeding (determined by confluency reaching 80-90%) differentiation was initiated by adding media A supplemented with Matrigel (D1) and continued as written in the manufacturer’s protocol. D16 cardiomyocytes were dissociated with STEMdiff Cardiomyocyte Dissociation Media and seeded for subsequent experiments as described below.

*Flow cytometry analysis of CMs*
CMs and iPSCs were prepared for flow cytometry as per Inside Stain Kit protocol (Miltenyi Biotec). Cells were stained with recombinant (REA400) FITC-conjugated anti-cardiac Troponin T (1:10) for 10 minutes. iPSCs were used as negative controls with unstained CMs acting as gating controls. Samples were analyzed on an LSR II (BD Biosciences).

*Calcium Imaging of CMs*	D16 contractile cardiomyocytes were dissociated and 6x10^5^ cells per well were seeded in Matrigel-coated 96-well plates in STEMdiff Cardiomyocyte Support Media (STEMCELL Technologies). The following day the media was replaced with STEMdiff Cardiomyocyte Maintenance Media and every two days thereafter. D34, cells were treated with 1 µM Fluo-4 AM (Invitrogen) in HBSS solution for 30 minutes at 37°C. Following Fluo-4 loading, cells were washed one time with HBSS solution. Imaging was carried out with a Nikon TE2000 microscope at 37°C with a 10x lens using Volocity software. Images were captured with 488 nm excitation and 516 nm emission at a rate of four images per second for 30 seconds. Image stacks were converted to videos with Volocity software (Quorum Technologies) and regions of interest were analyzed for changes in fluorescence intensity (f-f0/f0), with the resting fluorescence value F0 set as the minimal value recorded. Background intensity was subtracted from all values, and plots were normalized to zero fluorescence. Calcium transient amplitude and beat rate were calculated with Volocity software.

*xCELLigence RTCA data collection*
xCELLigence E-plates were coated with fibronectin in water for one hour prior to cell seeding. Before adding cells, a baseline recording was taken while only media was present in the well. 4x10^5^ dissociated CMs were seeded per well, then the plate was transferred to the device in a humidified incubator at 37^o^C, 5% CO_2_ and normoxic conditions. Twenty second sweeps were collected every three hours with a Cardio speed of 2.0 ms and ECR speed of 0.1 ms using RTCA CardioECR Software without stimulation (V1.2.0.1603; ACEA Biosciences). Recordings were paused every two days to change media to fresh maintenance media.

*CM statistical analyses*	Statistics for Ca^2+^ imaging or xCELLigence assays were calculated using RStudio. Normality was checked by ggqqplot and Shapiro.test. If data was normally distributed, analysis of variance was calculated followed by Tukey ANOVA followed by Tukey’s test. Where data was non-normal Kruskal-Wallis tests were used instead and pairwise comparisons made using Dunn’s test.

*Differentiation to hepatocytes*
All PGPC human iPSC lines were maintained in mTeSR1 as previously described. Prior to the induction of endoderm in the monolayer cultures, human iPSCs were passaged onto a Matrigel coated surface for 1 day in mTeSR1 based medium supplemented with 10 µM ROCK Inhibitor Y-27632. D1, differentiation towards definitive endoderm was carried out using STEMdiff Definitive Endoderm Kit (STEMCELL Technologies) for four days. D5, the resultant definitive endoderm (DE) cells in one of the 6-well-plate were reseeded in 12 wells of 24-well-plate, cultured in serum-free-differentiation (SFD)-based medium with B27 and 5 ng/ml basic fibroblast growth factor (bFGF) for two days and 100 ng/ml activin A for four days. The media was changed every other day. D8, DE was specified to foregut progenitor (FP) cells.
Bipotent hepatoblasts (HBs) were generated by culturing FPs in low glucose DMEM plus B27 supplemented with 40 ng/ml bFGF, 10 µM SB-431542 and 50 ng/ml bone morphogenic protein (BMP4) from day 8 to day 14, media changed every other day. To promote expansion and maturation of the hepatoblast population, the HBs were cultured in a mixture of low glucose DMEM / Ham’s F12 (3:1) media with 0.1% BSA, 1% vol/vol B27+retinoic acid (RA) supplement, ascorbic acid, glutamine, MTG, 20 ng/ml hepatocyte growth factor (HGF), 40 ng/ml dexamethasone (Dex) and 20 ng/ml oncostatin M (OSM) for 7 days. At D22, cells were cultured in a mixture of high glucose DMEM / Ham’s F12 (3:1) with 0.1% BSA, 1% vol/vol B27+RA supplement, ascorbic acid, glutamine, MTG, 20 ng/ml HGF, 40 ng/ml Dex and 20 ng/ml OSM for 4 days to generate hepatocyte like cells (HLCs) at D25. The media was changed every other day.

*Flow cytometry analysis during hepatocyte-like cell differentiation*
Cells were prepared for flow cytometry by dissociating to single cells with TrypLe, washed then fixed with 4% PFA. Cells were permeabilized with Perm/Wash buffer (BD Biosciences) then labeled with primary antibodies at concentrations listed in Table S8. Samples were analyzed on an LSR II (BD Biosciences).

*Cytochrome p450 activity*	CYP3A4/7 activity in HLCs was measured using the p450 Glo assay kit (Promega) according to the manufacturer’s instructions and a P450-Glomax 96 microplate luminometer (Promega). HBs were used as negative controls and 1 µM ketoconazole was used as an inhibitor of CYP3A4/3A7.

*Differentiation to kidney organoids*	Single-cell suspensions of iPSCs were made using Accutase and cells were seeded at densities of 7,500-12,000 cells/cm^2^ on freshly prepared Matrigel-coated plates with mTeSR1 containing 10 μM Y-27632. Differentiation was started (D0) by changing medium to APEL-2 supplemented with 8 μM CHIR99021. Medium was fully replaced every other day until D5 where CHIR-medium was switched to APEL-2 supplemented with 200 ng/ml human fibroblast growth factor (hFGF)-9 and 1 μg/ml heparin. iPSCs and differentiated cells were harvest for mRNA analysis at indicated time-points (D0, 4 and 7).
D7 cells were dissociated using Trypsin-EDTA for 1 minute at 37°C. Two milliliters of DMEM:F12 supplemented with 10% fetal bovine serum was added to inactivate Trypsin and cells were spun down at 300 g for 4 minutes. Supernatants were removed and cell pellets were resuspended in APEL-2 medium without differentiation factors. Aliquots of suspension containing 5x10^5^ cells were transferred to 1.5 ml microcentrifuge tubes and aggregated by centrifugation at 400 g for 3 minutes. Three to four aggregates were carefully transferred to 6-well Transwell cell culture plates using a wide-bore P200 pipette tip with minimal carry-over of medium. Initial Transwell medium consisted of APEL-2 with 5 μM CHIR99021. Medium was switched to APEL-2 with 200 ng/ml hFGF9 and 1 μg/ml heparin after one hour. Medium was replaced every other day thereafter. From D12 onward, APEL-2 was supplemented with heparin only.

*Immunostaining kidney organoids*	D25 organoids were fixed in 2% paraformaldehyde as described previously (Takasato et al., 2016). Organoids were embedded in paraffin and sectioned (6 μm sections) by the Core Pathology laboratory at The Centre for Phenogenomics (Toronto, ON, Canada). The immunofluorescence protocol was described previously (Rowan et al., 2018). In brief: sections were stained with the primary antibodies against Wilms Tumor 1 (WT1) (1:100) and E- Cadherin (ECAD (1:200) with appropriate secondary antibodies (Table S8). In addition, fluorescein-labeled Lotus Tetragonolobus lectin (LTL, 1:200) was used to stain the proximal tubule segments. LTL was added with secondary antibodies. After mounting, sections were imaged using an epifluorescence microscope.

*mRNA expression analysis of kidney organoid and HLC differentiations*	mRNA was isolated from iPSCs and differentiated cells using RNeasy Mini kit or TRIzol according to protocol (Qiagen). mRNA concentrations were determined using Nanodrop 2000C. mRNA was reverse transcribed using SuperScript First-Strand kit. Quantitative real-time PCR was performed using the ViiA7 and PowerSYBR Green PCR technology (kidney; ThermoFisher) or Advanced qPCR Mastermix with Supergreen Lo-Rox (HLC; Wisent). Sequences of primers used were either derived from previous publications or newly designed using Nucleotide Blast (https://blast.ncbi.nlm.nih.gov/Blast.cgi). Quantitation was performed using the standard curve method with serial dilutions of pooled cDNA of all samples. mRNA expression levels were normalized to ribosomal 18S or TBP expression and expressed relative to reference group as indicated.

*T-cell differentiation*
PGPC lines and the commercially available iPSC11 (Alstem Cell Advancements) were cultured on plates coated with Matrigel (Corning) with TeSR-E8 medium (STEMCELL Technologies). iPSCs were first differentiated to generate CD34+ hemogenic endothelial (HE) progenitor cells (as described previously in Kennedy et al., 2012), which were isolated by staining with anti-human CD34-PE (BD) and anti-PE-microbeads (Miltenyi) and enriched using magnetic-activated cell sorting (MACS) columns (Miltenyi). CD34+ cells were then co-cultured with OP9-DL4 cells to direct differentiation towards T-lineage cells as described previously in La Motte-Mohs, 2005. T-lineage cells were passaged onto new OP9-DL4 cells every 4-5 days and analyzed by flow cytometry using the following T-lineage markers: CD7, CD5, CD8a, CD1a, CD4, CD45, CD3, TCRαβ, TCRγδ, CD34 and CD8b. Flow cytometry data was collected on LSR-II (BD) and analyzed using FlowJo.

*Culture conditions and neural induction of sensory neurons*	Primary sensory neurons (PSNs) were generated using a protocol modified from Chambers et al., 2012. IPSCs (PGP17_11) were dissociated to single cells with Accutase and plated on Matrigel coated plates at 3.1x10^5^ - 3.7x10^5^ cells/cm^2^ in mTeSR1 + 10 µM Y-27632. Upon reaching confluence, neural differentiation was initiated (D0) by the addition of 100 nM LDN-193189 and 10 µM SB431542 in mTeSR1 media. On D2, media was replaced with mTeSR1 containing 100 nM LDN-193189, 10 µM SB431542, 3 µM CHIR99021, 10 µM SU5402, 10 µM DAPT, 10 ng/ml NGF and 1 µg/ml laminin. On D4, media was replaced with a 75% mTeSR1 / 25% N2 media (47.5% DMEM/F12, 47.5% neurobasal, 2% B-27 supplement, 1% N2 supplement, 1% Glutamax and 1% Pen/Strep) combination containing 100 nM LDN-193189, 10 µM SB431542, 3 µM CHIR99021, 10 µM SU5402, 10 µM DAPT, 10 ng/ml NGF and 1 µg/ml laminin. On D6, media was replaced with a 50% mTeSR1 / 50% N2 media combination containing 3 µM CHIR99021, 10 µM SU5402, 10 µM DAPT, 10 ng/ml NGF and 1 µg/ml laminin. On D8, media was replaced with a 25% mTeSR1 / 75% N2 media combination containing 3 µM CHIR99021, 10 µM SU5402, 10 µM DAPT, 10 ng/ml NGF and 1 µg/ml laminin. On D10, 100% N2 media was used containing 3 µM CHIR99021, 10 µM SU5402, 10 µM DAPT, 10 ng/ml NGF and 10 µM Ara-C. At D12, sensory neurons were dissociated with Accutase, washed with DMEM/F12 + 10% BSA and triturated until no visible clumps remained. The cell suspension was then filtered through a 70 µm filter, pelleted at 1100 rpm for 5 min and resuspended in N2 media containing 10 ng/ml NGF, 10 ng/ml BDNF, 10 ng/ml GDNF, 10 µg/ml laminin and 10 µM Y-27632. Neurons were reseeded onto plastic coverslips coated with 1 mg/ml polyornithine overnight followed by 10 µg/ml fibronectin and 40 µg/ml laminin also overnight at 2x10^5^ – 2.5x10^5^ cells/cm^2^ with media changed using the above combination twice weekly.

*Patch-clamp recordings*	Neurons were recorded in whole cell configuration with 70% series resistance compensation using an Axopatch 200B (Molecular Devices, Sunnyvale, CA, USA). Patch clamp recordings were performed at room temperature with traces acquired at 25 kHz and low-pass filtered at 2 kHz. Data was digitized with a power1401-3A data acquisition interface (CED ltd., Cambridge, England). Artificial cerebrospinal fluid (aCSF) contained the following in (mM): 126 NaCl, 2.5 KCl, 2 CaCl_2_, 2 MgCl_2_, 10 D-Glucose, 26 NaHCO_3_, 1.25 NaH_2_PO_4_ bubbled with 95% oxygen and 5% carbon dioxide, and was continuously perfused over the coverslips at a rate of 2 ml/min. Borosilicate glass microelectrodes (World Precision Instruments, Sarasota, FL USA) with a tip resistance of 4-6 MΩ were filled with solution containing the following (in mM): 125 KMeSO_4_, 5 KCl, 10 HEPES, 2 MgCl_2_, 4 Na_2_ATP and 0.4 Tris-GTP (ThermoFisher Scientific); pH was adjusted with KOH to 7.2. All membrane potential values were corrected for a junction potential of 9 mV. Rheobase and spiking patterns were assessed at a holding potential of -65mV. Repetitive spiking neurons were defined as those that responded to a supra-threshold current injection with 3 or more spikes. Cells which showed a resistance change greater than 20% over the course of the recording were rejected from subsequent analysis.

*Ca^2+^ imaging and analysis of PSNs*	Ca^2+^ imaging was performed on neurons incubated in 5 µM Ca^2+^ green-1 AM dye (ThermoFisher Scientific, Waltham, MA, USA) at room temperature for 20 minutes in oxygenated, HEPES buffered ACSF. Images were acquired using a NeuroCCD-SM256 imaging system (RedShirt Imaging, Decatur, GA) mounted on a Zeiss Examiner A1 with dichroic filter set 46HE. Images were acquired at a rate of 25 Hz with a 10X digital gain. Image analysis was performed by drawing a region of interest around individual neurons using Neuroplex software and exporting the raw fluorescence values to SigmaPlot for subsequent analysis. Healthy cells were identified by application of 500mM KCl using a Toohey Spritzer pressure system IIe fluid delivery system (Fairfield, NJ) at 5 PSI with a 4-5 second pulses. The tip of the puff pipette was positioned 50-70 µm from the cell group and 5 µm above the coverslip surface while undergoing Ca^2+^ imaging. Neurons that showed a robust rise in cytosolic Ca^2+^ to KCl puffs were considered healthy. Phenotyping was performed on these cells with the same picospritzer settings using 3 mM GABA, 20 µM capsaicin and 200 µM ATP in HEPES buffered ACSF. To differentiate ionotropic from metabotropic GABA receptors, 100 µM picrotoxin was bath applied following the initial identification of GABA responsive cells. All drugs were acquired from Sigma-Aldrich.

*Statistical analysis of PSNs*
Comparisons between time points and spiking patterns were conducted using Mann-Whitney U ranked sum tests. Chi Square tests were performed to assess developmental differences in agonist sensitivities and spiking patterns.

*Primer list*

| Target | Forward primer | Reverse primer |
| --- | --- | --- |
| Sendai virus detection | GGATCACTAGGTGATATCGAGC | ACCAGACAAGAGTTTAAGAGATATGTATC |
| *AR* | (FAM)CGTGCGCGAAGTGATCCAGA | GTTTTTTGCTGCTGCCTGGGGCTAGT |
| MYBPC3 CRISPR/Cas9 targeting | CACCGGACTCCTGCACAGTACAGT | AAACACTGTACTGTGCAGGAGTCC |
| MYBPC3 (gDNA) | CTGTGGCGGTTAGTTGGAGT | GTAGACGCGCATCTCGTACA |
| *TBP* | TAAGAGAGCCACGAACCACG | TTCACATCACAGTCCCCAC |
| *GATA6* | GCCACTACCTGTGCAACGCCT | CAATCCAAGCCGCCGTGATGAA |
| *FOXA2* | ATGCACTCGGCTTCCAGTATG | TGTTGCTCACGGAAGAGTAGC |
| *HNF6* | GTGTTGCCTCTATCCTTCCCAT | CGCTCCGCTTAGCAGCAT |
| *HNF4A* | CATGGCCAAGATTGACAACCT | TTCCCATATGTTCCTGCATCAG |
| *CYP3A7* | GACGGGCTTCATCCAATGTG | TGGGGGTGGTGGAGATAGTC |
| *CEBPA* | GGAGCTGAGATCCCGACAAG | GCGTGGAACTAGAGACCCTC |
| *ALB* | CCTTTGGCACAATGAAGTGGGTAACC | CAGCAGTCAGCCATTTCACCATAG |
| *AFP* | AGAACCTGTCACAAGCTGTG | GACAGCAAGCTGAGGATGTC |
| *A1AT* | ACATGTGAGCACGGAGAACA | TTCTGGGACACTAGAGTCGTG |
| *18S* | GATGGGCGGCGGAAAATAG | GCGTGGATTCTGCATAATGGT |
| *GATA3* | GCCCCTCATTAAGCCCAAG | TTGTGGTGGTCTGACAGTTCG |
| *HOXD11* | GCCAGTGTGCTGTCGTTCCC | CTTCCTACAGACCCCGCCGT |
| *T* | AGGTACCCAACCCTGAGGA | GCAGGTGAGTTGTCAGAATAGGT |
| *NPHS1* | TATCCCGCCCAGAAACTGTG | CGCGAGTCCTTGTACCACAT |
| *SLC3A1* | TCGCTCAAAGTCACCAATGC | GCTGAGTCTTTTGGACATCAACAT |
| *SLC12A1* | GGATGGGTGAAAGGTGTGCT | GAACTCCAAGACCAATTCCAGC |
| *PECAM* | CATTACGGTCACAATGACGATGT | ATCTGCTTTCCACGGCATCA |
| *FOXD1* | TTGGGGACTCTGCACCAAGG | TTCCTCGAGCGCGCTAACAT |

*Antibody and cell labeling reagent list*

| Target | Species | Concentration (Technique) | Company | Catalogue number |
| --- | --- | --- | --- | --- |
| SSEA4 | Mouse | 1:100 (ICC) | ThermoFisher Scientific | 41-4000 |
| TRA1-60 | Mouse | 1:100 (ICC) | ThermoFisher Scientific | 41-1000 |
| NANOG | Rabbit | 1:400 (ICC) | Cell Signaling Technology | 4903 |
| OCT4 | Rabbit | 1:200 (ICC) | Abcam | ab19857 |
| TUBB3 | Mouse | 1:200 (ICC) | Milipore Sigma | MAB1637 |
| SMA | Mouse | 1:200 (ICC) | ThermoFisher Scientific | 18-0106 |
| AFP | Mouse | 1:200 (ICC) | R&D Systems | MAB1368 |
| MAP2 | Guinea pig | 1:1000 (ICC) | Synaptic Systems | 188 004 |
| GFP | Chicken | 1:500 (ICC) | ThermoFisher Scientific | A-10262 |
| cTNT | Mouse | 1:4000 (ICC) | Abcam | ab 8295 |
| cTNT-FITC conjugate | Recombinant human | 1:10 (Flow) | Miltenyi Biotec | 130-106-687 |
| MLC2V | Rabbit | 1:400 (ICC) | Abcam | Ab79935 |
| MYBPC3 | Rabbit | 1:1000 (WB) | Abcam | ab171153 |
| Actinin | Mouse | 1:200 (ICC) | Abcam | Ab9465 |
| ALB | Rabbit | 1:50 (ICC/Flow) | DAKO | F0117 |
| Alpha-1-AT | Goat | 1:100 (Flow) | Bethyl | A80-122A |
| AFP-APC conjugate | Mouse | 1:200 (Flow) | Sino Biological Inc. | 121177-MM35-A |
| CYP3A7 | Mouse | 1:100 (ICC/Flow) | NOVUS | BP2-37502SS |
| HNF4A | Rabbit | 1:100 (ICC) | Cell Signaling | 3113S |
| WT1 | Mouse | 1:100 (IHC) | DAKO | M3561 |
| ECAD | Rabbit | 1:200 (IHC) | Cell Signaling Technology | 3195 |
| Lotus Tetragonolobus lectin (Fluorescein-labeled) | N/A | 1:200 (IHC) | Vector Laboratories | FL-1321 |
| Anti-mouse IgG (AlexaFluor 555) | Goat | 1:1000 (ICC) | ThermoFisher Scientific | A21422 |
| Anti-mouse IgM (AlexaFluor 555) | Goat | 1:1000 (ICC) | ThermoFisher Scientific | A21426 |
| Anti-rabbit (AlexaFluor 488) | Goat | 1:1000 (ICC) | ThermoFisher Scientific | A11008 |
| Anti-mouse (AlexaFluor 488) | Goat | 1:1000 (IHC) | ThermoFisher Scientific | A21042 |
| Anti-mouse (AlexaFluor 405) | Goat | 1:1000 (IHC) | ThermoFisher Scientific | A31553 |
| Anti-rabbit (AlexaFluor 568) | Donkey | 1:1000 (IHC) | ThermoFisher Scientific | A10042 |
| IRDye 800CW anti-rabbit | Goat | 1:20000 (WB) | LI-COR Biosciences | 925-32211 |
| CD7-FITC |  | 1:100 (Flow) | BioLegend | 343104 |
| CD5-PerCP/Cy5.5 |  | 1:100 (Flow) | BioLegend | 300620 |
| CD8a-PE/CF594 |  | 1:100 (Flow) | BioLegend | 301058 |
| CD1a-APC |  | 1:100 (Flow) | BioLegend | 300110 |
| CD4-AF700 |  | 1:100 (Flow) | BioLegend | 300526 |
| CD45-APC/Cy7 |  | 1:100 (Flow) | BioLegend | 304014 |
| CD3-BV510 |  | 1:100 (Flow) | BioLegend | 317332 |
| TCRαβ-APC |  | 1:100 (Flow) | BioLegend | 306718 |
| TCRγδ-FITC |  | 1:100 (Flow) | BioLegend | 331208 |
| CD34-PE |  | 1:25 (Flow) | BD | 550761 |
| CD8b-PE/Cy7 |  | 1:100 (Flow) | eBiosciences | 25-5273-42 |

Abyzov, A., Urban, A.E., Snyder, M., and Gerstein, M. (2011). CNVnator: an approach to discover, genotype, and characterize typical and atypical CNVs from family and population genome sequencing. Genome Res. *21*, 974–984.

Van der Auwera, G.A., Carneiro, M.O., Hartl, C., Poplin, R., del Angel, G., Levy-Moonshine, A., Jordan, T., Shakir, K., Roazen, D., Thibault, J., et al. (2013). From FastQ Data to High-Confidence Variant Calls: The Genome Analysis Toolkit Best Practices Pipeline. Curr. Protoc. Bioinforma. *43*, 11.10.1-11.10.33.

Benchling (2019). Benchling [Biology Software].

Chambers, S.M., Qi, Y., Mica, Y., Lee, G., Zhang, X.J., Niu, L., Bilsland, J., Cao, L., Stevens, E., Whiting, P., et al. (2012). Combined small-molecule inhibition accelerates developmental timing and converts human pluripotent stem cells into nociceptors. Nat. Biotechnol. *30*, 715–720.

Cibulskis, K., Lawrence, M.S., Carter, S.L., Sivachenko, A., Jaffe, D., Sougnez, C., Gabriel, S., Meyerson, M., Lander, E.S., and Getz, G. (2013). Sensitive detection of somatic point mutations in impure and heterogeneous cancer samples. Nat. Biotechnol. *31*, 213–219.

DePristo, M.A., Banks, E., Poplin, R., Garimella, K. V, Maguire, J.R., Hartl, C., Philippakis, A.A., del Angel, G., Rivas, M.A., Hanna, M., et al. (2011). A framework for variation discovery and genotyping using next-generation DNA sequencing data. Nat. Genet. *43*, 491–498.

Hotta, A., Cheung, A.Y.L., Farra, N., Garcha, K., Chang, W.Y., Pasceri, P., Stanford, W.L., and Ellis, J. (2009). EOS lentiviral vector selection system for human induced pluripotent stem cells. Nat. Protoc. *4*, 1828–1844.

Kennedy, M., Awong, G., Sturgeon, C.M., Ditadi, A., LaMotte-Mohs, R., Zúñiga-Pflücker, J.C., and Keller, G. (2012). T lymphocyte potential marks the emergence of definitive hematopoietic progenitors in human pluripotent stem cell differentiation cultures. Cell Rep. *2*, 1722–1735.

Li, H., and Durbin, R. (2009). Fast and accurate short read alignment with Burrows-Wheeler transform. Bioinformatics *25*, 1754–1760.

La Motte-Mohs, R.N. (2005). Induction of T-cell development from human cord blood hematopoietic stem cells by Delta-like 1 in vitro. Blood *105*, 1431–1439.

Reuter, M.S., Walker, S., Thiruvahindrapuram, B., Whitney, J., Cohn, I., Sondheimer, N., Yuen, R.K.C., Trost, B., Paton, T.A., Pereira, S.L., et al. (2018). The Personal Genome Project Canada: findings from whole genome sequences of the inaugural 56 participants. Can. Med. Assoc. J. *190*, E126–E136.

Rowan, C.J., Li, W., Martirosyan, H., Erwood, S., Hu, D., Kim, Y.-K., Sheybani-Deloui, S., Mulder, J., Blake, J., Chen, L., et al. (2018). Hedgehog-GLI signaling in Foxd1- positive stromal cells promotes murine nephrogenesis via TGFβ signaling. Development *145*, 1–13.

Takasato, M., Er, P.X., Chiu, H.S., and Little, M.H. (2016). Generation of kidney organoids from human pluripotent stem cells. Nat. Protoc. *11*, 1681–1692.

Wang, K., Li, M., and Hakonarson, H. (2010). ANNOVAR: functional annotation of genetic variants from high-throughput sequencing data. Nucleic Acids Res. *38*, e164–e164.

Yuen, R.K.C., Thiruvahindrapuram, B., Merico, D., Walker, S., Tammimies, K., Hoang, N., Chrysler, C., Nalpathamkalam, T., Pellecchia, G., Liu, Y., et al. (2015). Whole-genome sequencing of quartet families with autism spectrum disorder. Nat. Med. *21*, 185–191.

Zhu, M., Need, A.C., Han, Y., Ge, D., Maia, J.M., Zhu, Q., Heinzen, E.L., Cirulli, E.T., Pelak, K., He, M., et al. (2012). Using ERDS to infer copy-number variants in high-coverage genomes. Am. J. Hum. Genet. *91*, 408–421.

Table S1. Summary of PGPC iPSC characterizations.

| Cell line | Gender | Age | Mycoplasma | Sendai virus | Karyotype | X-chromosome active | Pluripotency staining | Embryoid body assay | Additional information |
| --- | --- | --- | --- | --- | --- | --- | --- | --- | --- |
| PGPC3_23 | M | 48 | negative | negative | normal | N/A | Oct4, Nanog, SSEA4, Tra-1-60 positive | TUBB3, SMA, AFP positive |  |
| PGPC3_66 | M | 48 | negative | negative | normal | N/A | Oct4, Nanog, SSEA4, Tra-1-60 positive | TUBB3, SMA, AFP positive |  |
| PGPC3_75 | M | 48 | negative | negative | normal | N/A | Oct4, Nanog, SSEA4, Tra-1-60 positive | TUBB3, SMA, AFP positive | enhanced characterization |
| PGPC3_93 | M | 48 | negative | negative | normal | N/A | Oct4, Nanog, SSEA4, Tra-1-60 positive | TUBB3, SMA, AFP positive |  |
| PGPC3_87 | M | 48 | negative | negative | normal | N/A | Oct4, Nanog, SSEA4, Tra-1-60 positive | TUBB3, SMA, AFP positive |  |
| PGPC14_26 | F | 62 | negative | negative | normal | Skewed—X_1_ | Oct4, Nanog, SSEA4, Tra-1-60 positive | TUBB3, SMA, AFP positive | enhanced characterization |
| PGPC14_70 | F | 62 | negative | negative | normal | Skewed—X_1_ | Oct4, Nanog, SSEA4, Tra-1-60 positive | TUBB3, SMA, AFP positive |  |
| PGPC14_16 | F | 62 | negative | negative | normal | Skewed—X_1_ | Oct4, Nanog, SSEA4, Tra-1-60 positive | TUBB3, SMA, AFP positive |  |
| PGPC14_94 | F | 62 | negative | negative | normal | Skewed—X_1_ | Oct4, Nanog, SSEA4, Tra-1-60 positive | TUBB3, SMA, AFP positive |  |
| PGPC17_4 | M | 61 | negative | negative | normal | N/A | Oct4, Nanog, SSEA4, Tra-1-60 positive | TUBB3, SMA, AFP positive |  |
| PGPC17_11 | M | 61 | negative | negative | normal | N/A | Oct4, Nanog, SSEA4, Tra-1-60 positive | TUBB3, SMA, AFP positive | enhanced characterization |
| PGPC17_45 | M | 61 | negative | negative | normal | N/A | Oct4, Nanog, SSEA4, Tra-1-60 positive | TUBB3, SMA, AFP positive |  |
| PGPC17_80 | M | 61 | negative | negative | normal | N/A | Oct4, Nanog, SSEA4, Tra-1-60 positive | TUBB3, SMA, AFP positive |  |
| PGPC17_142 | M | 61 | negative | negative | normal | N/A | Oct4, Nanog, SSEA4, Tra-1-60 positive | TUBB3, SMA, AFP positive |  |
| PGPC1_30 | F | 46 | negative | negative | normal | Skewed—X_1_ | Oct4, Nanog, SSEA4, Tra-1-60 positive | TUBB3, SMA, AFP positive |  |
| PGPC1_67 | F | 46 | negative | negative | normal | Skewed—X_2_ | Oct4, Nanog, SSEA4, Tra-1-60 positive | TUBB3, SMA, AFP positive |  |
| PGPC1_73 | F | 46 | negative | negative | normal | Skewed—X_2_ | Oct4, Nanog, SSEA4, Tra-1-60 positive | TUBB3, SMA, AFP positive |  |
| PGPC1_96 | F | 46 | negative | negative | normal | Skewed—X_1_ | Oct4, Nanog, SSEA4, Tra-1-60 positive | TUBB3, SMA, AFP positive |  |

Table S2 (excel). Comprehensive list of genomic variants in PGPC donor and iPSC lines.

Table S3. Genomic coding variants of concern in PGPC donors/cell lines and publicly available iPSC lines.

|  | **Gene, accession number** | **Variant, zygosity** | **Disease/biological function, inheritance** | **Interpretation** |
| --- | --- | --- | --- | --- |
| **PGPC1** | N/A | N/A | N/A | N/A |
| **PGPC3** | *TRPM4*, NM_017636 | c.2531G>A, p.(Gly844Asp), het | Progressive familial heart block, AD | VUS |
| **PGPC3_75** | *ZNF283*, NM_001297752 | c.63C>A, p.(Cys21*), het | Unknown. | Uncertain, pLI=0 |
| **PGPC14** | *KCNE2*, NM_172201 | c.29C>T, p.(Thr10Met), het | Long QT syndrome; Atrial fibrillation, AD | VUS |
| **PGPC14_26** | *TRIM71*, NM_001039111 | c.1963C>T, p.(Arg655*), het | Ubiquitin ligase. Neural differentiation, brain development. Embryonic stem cell proliferation. | Uncertain, pLI=0.99 |
| **PGPC14_26** | *FRMD4A*, NM_001318337 | c.563+1G>T, het | Cell polarity, associated with Alzheimer’s disease risk. | Uncertain, pLI=1.0 |
| **PGPC14_26** | *IL1RAPL1* | chrX:29319382-29335268 intronic deletion, het | Presynaptic differentiation during synapse formation, associated with nonsyndromic X-linked recessive intellectual disability | VUS |
| **PGPC17** | N/A | N/A | N/A | N/A |
| **PGPC17_11** | *ROBO2*, NM_001290039 | c.2621G>A, p.(Trp874*), het | Slit receptor. Axon branching, dendritic patterning, and neuronal migration. Endothelial cell migration and angiogenesis. Heart tube formation. Myotome formation. | Uncertain, pLI=1.0 |
| **HPSI0114i.kolf_2** | *COL3A1*, NM_000090 | c.[3526-1G>A; 3526G>A], p.[?; (Gly1176Ser)], het | Ehlers-Danlos syndrome, type IV, AD | Likely pathogenic |
| **HPSI0114i.kolf_2** | *ARID2*, NM_152641 | c.590_608delCTAAAATCATCACTTTACT, p.(Pro197Hisfs*12), het | Coffin-Siris syndrome, AD | Likely pathogenic |
| **HPSI0114i-kolf_2** | *ZNF398*, NM_170686 | c.1420C>T, p.(Gln474*), het | Unknown. | Uncertain, pLI=0.96 |
| **HPSI0114i-kolf_2** | *UBE3C*, NM_014671 | c.661_668del TCAAGTAT, p.(Ser221*), het | Ubiquitin ligase. Cell proliferation. Wnt/β-catenin signaling. | Uncertain, pLI=1.0 |
| **HPSI0114i-kolf_2** | *CDC37*, NM_007065 | c.192C>A, p.(Cys64*), het | Cell cycle progression. | Uncertain, pLI=0.96 |
| **HPSI0214i-wibj_2** | N/A | N/A | N/A | N/A |
| **HPSI0314i.sojd_3** | *BCOR*, NM_017745 | c.1042C>T, p.(Gln348*), het | Oculofaciocardiodental syndrome, XLD | Likely pathogenic |
| **HPSI0314i-hoik_1** | *PTK2*, NM_001352736 | c.2551C>T, p.(Gln851*), het | Cell growth. Integrin-mediated signal transduction. Heart and blood vessel development. | Uncertain, pLI=1.0 |
| **HPSI0214i-kucg_2** | *TNS3*, NM_022748 | c.1239delC, p.(Arg414Glyfs*4), het | Cell migration. Focal adhesion. | Uncertain, pLI=1.0 |

Abbreviations: AD, autosomal dominant; het, heterozygous; PGPC, Personal Genome Project Canada; pLI, probability of loss-of-function intolerance (http://exac.broadinstitute.org/); VUS, variant of uncertain significance; XLD, X-linked dominant.

Table S4. Genome sequencing quality metrics of donor material, newly derived and publicly available iPSC lines.

| **Sample** | **Total reads** | **Median read depth** | **Region covered 10X [%]** | **Region covered 20X [%]** | **Region covered 30X [%]** |
| --- | --- | --- | --- | --- | --- |
| PGPC1 (donor) | 933,877,478 | 38 | 96.8 | 95.6 | 84.4 |
| PGPC3 (donor) | 963,792,408 | 39 | 97.3 | 93.7 | 84.5 |
| PGPC3_75 | 780,473,626 | 31 | 96.8 | 88.9 | 55.2 |
| PGPC14 (donor) | 958,002,700 | 39 | 97.0 | 95.9 | 87.0 |
| PGPC14_26 | 774,673,256 | 31 | 96.9 | 92.4 | 56.4 |
| PGPC17 (donor) | 779,923,976 | 33 | 96.8 | 90.8 | 64.7 |
| PGPC17_11 | 765,097,084 | 31 | 96.8 | 89.0 | 56.3 |
| PGPC17_11_MYBPC3_KO | 914,771,348 | 36 | 97.4 | 92.9 | 77.9 |
| HPSI0114i-kolf_2 | 785,913,438 | 31 | 96.1 | 89.0 | 57.9 |
| HPSI0214i-kucg_2 | 856,258,780 | 37 | 96.9 | 92.7 | 77.7 |
| HPSI0214i-wibj_2 | 841,290,742 | 35 | 97.0 | 95.2 | 74.8 |
| HPSI0314i-hoik_1 | 945,070,686 | 36 | 97.0 | 95.6 | 80.9 |
| HPSI0314i-sojd_3 | 964,602,272 | 32 | 96.7 | 94.0 | 63.4 |

Table S5 (excel). Comprehensive list of genomic variants in HipSci cell lines

Supplemental Figure 1

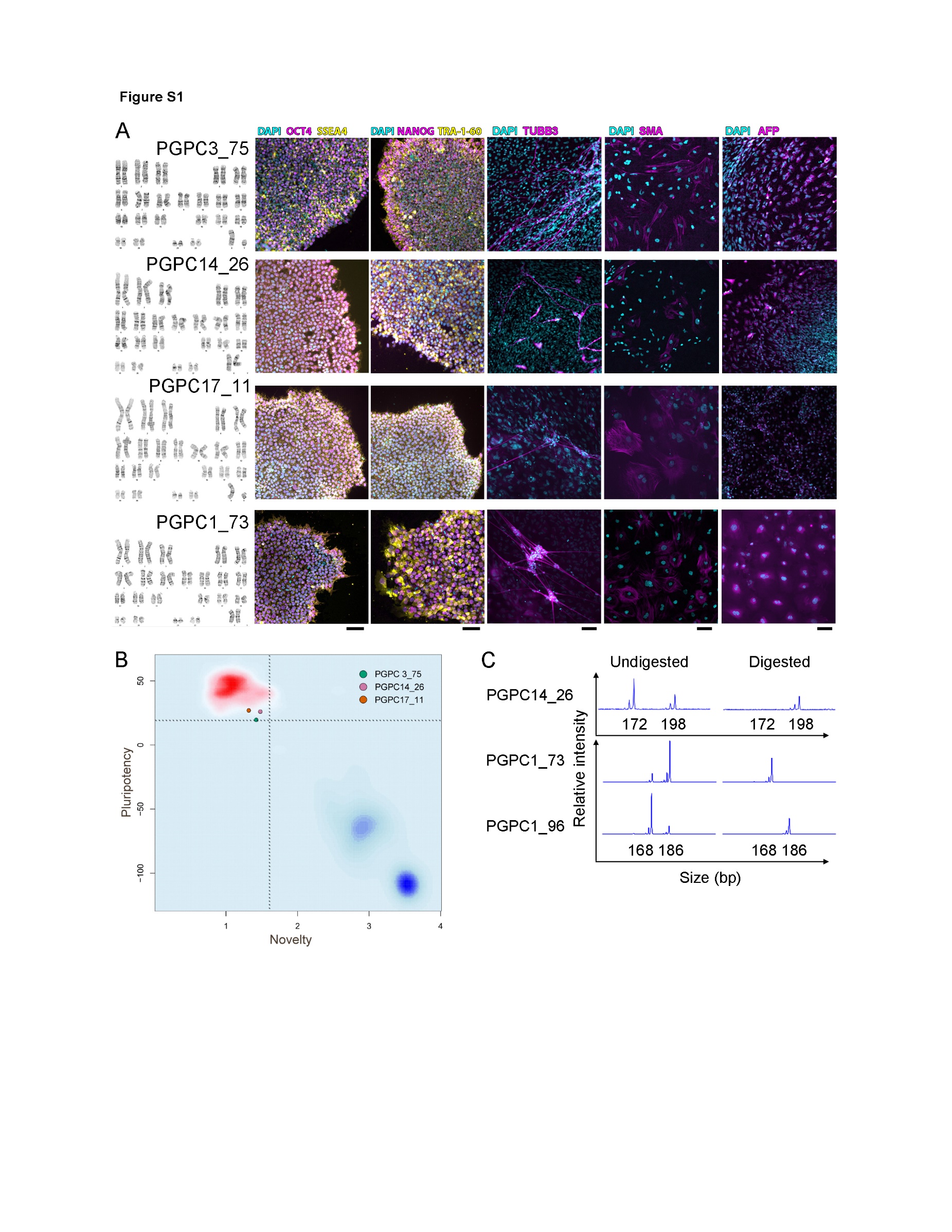

Figure S1. Quality control and pluripotency characterization of PGPC lines. Related to Table S1.
(A) Normal karyotype, positive pluripotency marker immunocytochemistry labeling (OCT4, SSEA4, NANOG, and TRA-1-60) of iPSC colonies, and germ layer marker labeling of embryoid body cultures (TUBB3, SMA, and AFP) were found in all PGPC cell lines. (B) PluriTest plot of PGPC3_75, 14_26, and 17_11 RNAseq data localized all three lines sequenced to date to the pluripotency quadrant. (C) Female iPSC lines were assessed for X chromosome inactivation via the androgen receptor assay. All lines showed clear separation between the two amplicons in the undigested control. After digestion with methylation-sensitive enzymes all lines show complete skewing towards either active X-chromosome as shown by the presence of a single peak cluster. Full skewing list found in Table S1.

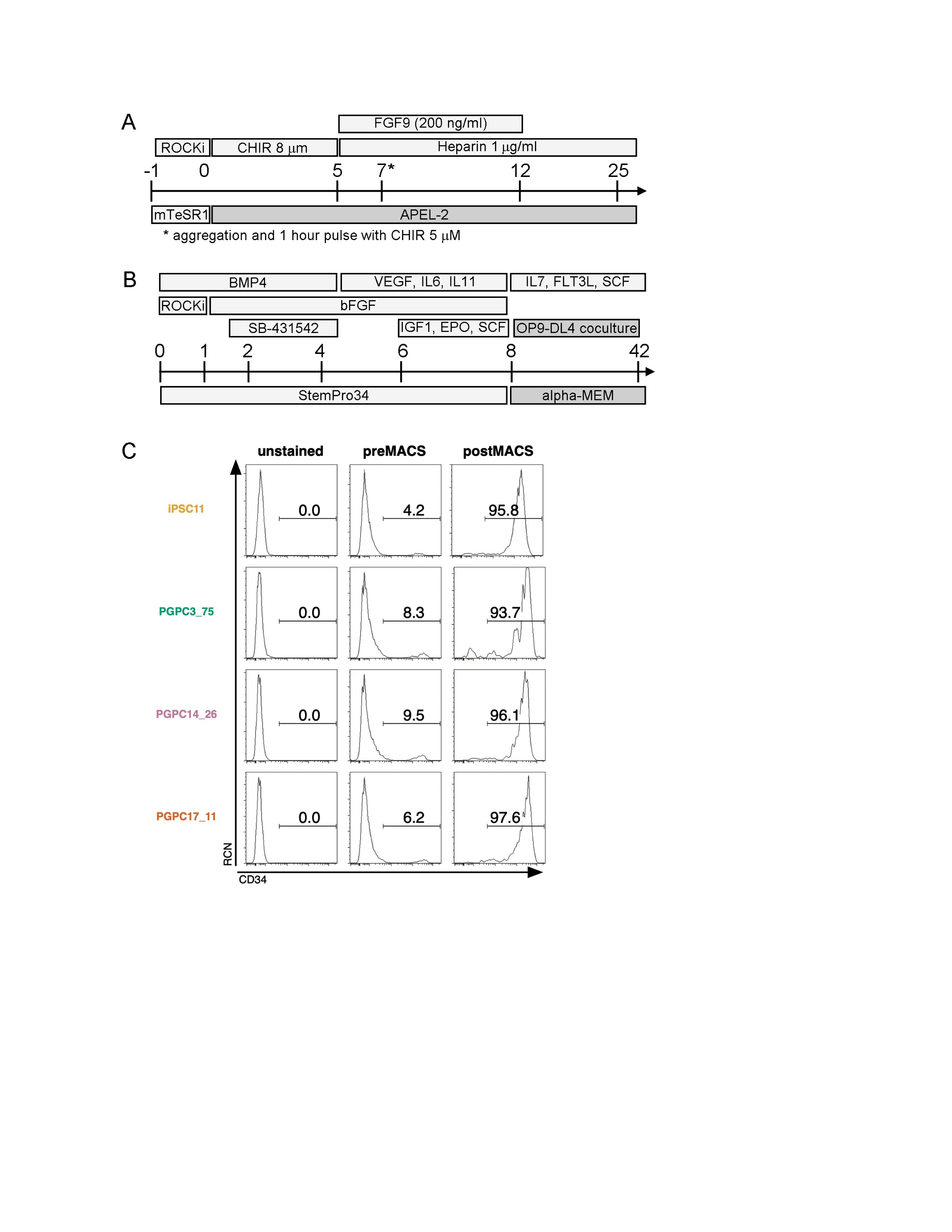
Supplemental Figure 2.

Figure S2. PGPC differentiation to kidney and T-cells. Related to Figure 4.
(A) Kidney organoid differentiation scheme. Single-cell suspensions of iPSCs were dissociated to single cells and grown in ROCK inhibitor for 24 hours before beginning differentiation by changing medium to APEL-2 supplemented with CHIR. Medium was fully replaced every other day until D5 where CHIR-medium was switched to APEL-2 supplemented with human fibroblast growth factor (hFGF)-9 and heparin. D7 cells were dissociated and aggregated by centrifugation in 1.5 ml tubes. Three to four aggregates were carefully transferred to Transwell cell culture plates and grown in media consisted of APEL-2 with CHIR. Medium was switched to APEL-2 with hFGF9 and heparin after one hour. Media was replaced every other day thereafter. From D12 onward, APEL-2 was only supplemented with heparin. (B) T-cell differentiation scheme. iPSCs were lifted with ReLeSR to clusters of 5-10 cells and transferred to low-cluster plates in StemPro34 media containing BMP4 and ROCK inhibitor. Media was changed after 24 hours to StemPro34 containing BMP4, CHIR and bFGF. At 42 hours, media was changed again now including SB-431542. At D4 EBs have a crumpled appearance and media was changed to include bFGF, VEGF, IL-6, IL-11. At D6 media was added on top of current culture media and contained VEGF, EPO, IGF, SCF, IL-6, IL-11, and bFGF. At D8, EBs were dissociated and enriched for CD34+ cells by MACS then co-cultured with OP9-DL4 cells in Alpha MEM containing Flt3-L, SCF and IL-7. Differentiating T-lineage cells were transferred to fresh OP9-DL4 cells approximately every 4-5 days. (C) Expression of CD34 measured by flow cytometry before and after MACS enrichment for CD34+ cells.

Supplemental Figure 3

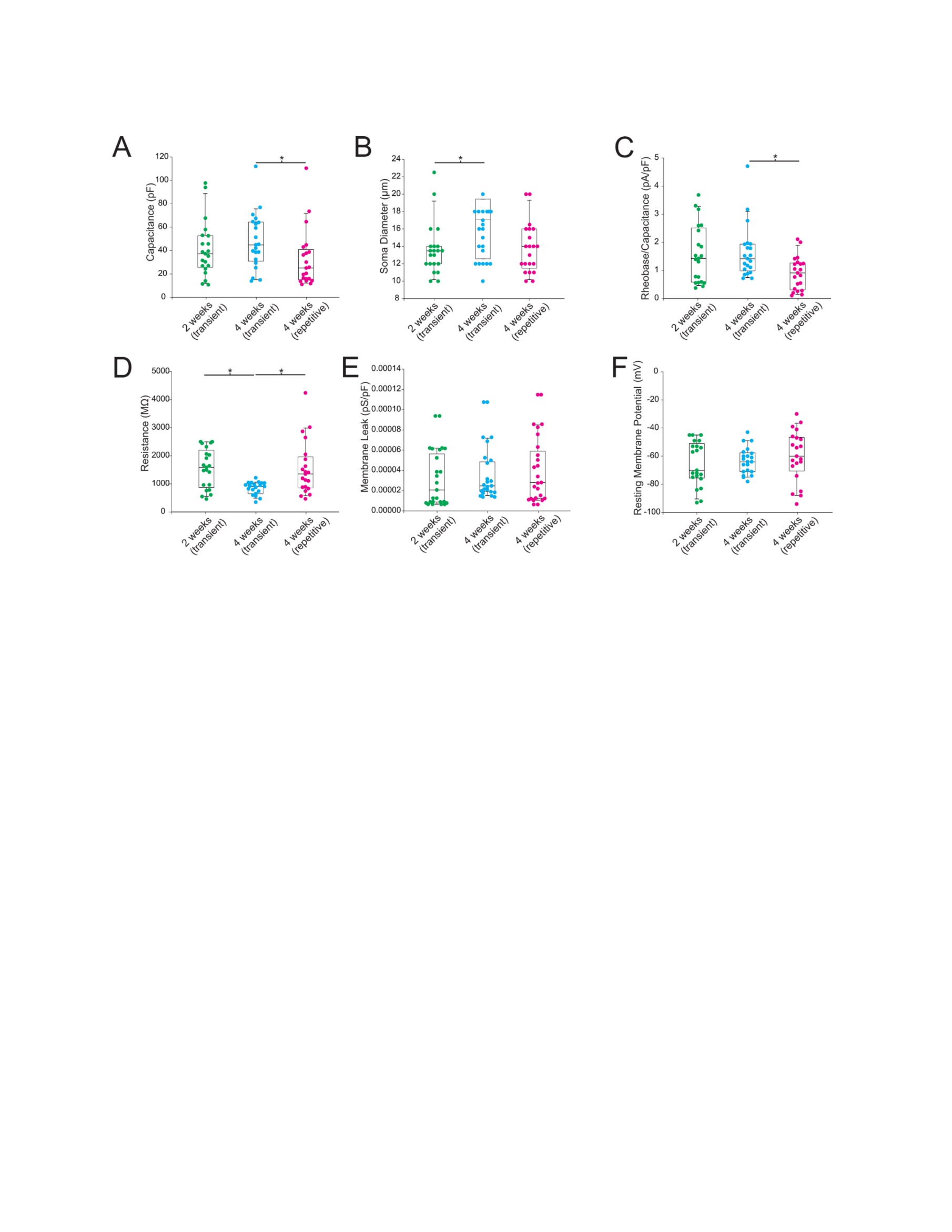

Figure S3. Comparison of other electrophysiological properties of sensory neurons. Related to Figure 5.
(A) membrane capacitance, which is proportional to membrane surface area, (B) soma diameter, (C) rheobase normalized by membrane capacitance, (D) input resistance, (E) membrane leak, based on the reciprocal of input resistance normalized by capacitance, and (F) resting membrane potential. * shows p < 0.05 based on Mann-Whitney U tests. Notably, differences in input resistance (D) were lost after normalizing by capacitance (A), which suggests that differences in input resistance were due primarily to cell size rather than membrane leakiness.
